# Primer biases in the molecular assessment of diet in multiple insectivorous mammals

**DOI:** 10.1101/2021.01.18.426998

**Authors:** Samuel S. Browett, Thomas G. Curran, Denise B. O’Meara, Andrew P. Harrington, Naiara Guimarães Sales, Rachael E. Antwis, David O’Neill, Allan D. McDevitt

## Abstract

Our understanding of trophic interactions of small insectivorous mammals has been drastically improved with the advent of DNA metabarcoding. The technique has continued to be optimised over the years, with primer choice repeatedly being a vital factor for dietary inferences. However, the majority of dietary studies examining the effect of primer choice often rely on *in silico* analyses or comparing single-niche species. Here we apply DNA metabarcoding to empirically compare the prey detection capabilities of two widely used primer sets when assessing the diets of a flying (lesser horseshoe bat; *Rhinolophus hipposideros*) and two ground dwelling insectivores (greater white-toothed shrew; *Crocidura russula* and pygmy shrew; *Sorex minutus*). Although *R. hipposideros* primarily rely on two prey orders (Lepidoptera and Diptera), the unique taxa detected by each primer shows that a combination of primers may be the best approach to fully describe bat trophic ecology. However, random forest classifier analysis suggest that one highly degenerate primer set detected the majority of both shrews’ diet despite higher levels of host amplification. The wide range of prey consumed by ground-dwelling insectivores can therefore be accurately documented from using a single broad-range primer set, which can decrease cost and labour. The results presented here show that dietary inferences will differ depending on the primer or primer combination used for insectivores occupying different niches (i.e. hunting in the air or ground) and demonstrate the importance of performing empirical pilot studies for novel study systems.

## Introduction

In a constantly changing environment, knowledge of complex food webs is vital for our understanding of ecosystem functioning and biodiversity conservation. The advent of Next-Generation Sequencing (NGS) technology has revolutionised the analyses of trophic interactions (Deagle et al. 2019; Browett et al. 2020), with DNA metabarcoding (i.e. simultaneous identification of multiple species using a standardised region of DNA) of faecal samples or gut contents becoming widely adopted for describing diets (Pompanon et al. 2012). Despite the significant developments and improvements afforded by DNA metabarcoding for dietary studies over the last decade, the technique has certain limitations. These include problems in describing diverse diets (e.g. omnivorous species); assigning sequences to appropriate taxonomic levels with incomplete or poor reference databases; false negatives/positives for species detections, and host co-amplification (Piñol et al. 2015; Alberdi et al. 2019; Deagle et al. 2019).

Several of these limitations are particularly evident when studying the diets of mammalian insectivores in terrestrial environments. Invertebrates are massively diverse and widely distributed (Stork 2018), which makes describing invertebrate-based diets via DNA metabarcoding challenging. Given that insectivores can potentially have a broad diet (Brown et al. 2014), a key consideration is the choice of primers to use due to varying detection capabilities (Corse et al. 2019), or target only specific invertebrate groups (Saitoh et al. 2016). To capture the expected wide range of invertebrate taxonomic groups, highly degenerative (non-specific) primers can be used, but studies comparing their efficiency have largely been restricted to analyses performed *in silico* (Piñol et al. 2018) or using bulk samples and/or mock communities (Elbrecht et al. 2019). While these are essential steps in primer design and have led to the ability to detect a wide range of invertebrate species, they may not account for some of the potential biases within a dietary context (i.e. predator/host amplification; Zeale et al. 2011). The broader the taxonomic range of the primers, the more likely the chance of amplifying non-target taxa and reducing the amount of information on a species diet.

In terms of insectivorous mammalian predators, bats are well-represented in dietary DNA metabarcoding studies due to their ecological importance and their significant role in the suppression of insects e.g. pests and vectors implicated in the spread of disease that may negatively impact agriculture (Galan et al. 2018; Baroja et al. 2019). They have not only served as a key study group for primer comparisons, but also for methodological development such as sampling design, evaluation of setting clustering thresholds for Molecular Operational Taxonomic Unit (MOTU), and mitigating contamination/errors (Alberdi et al. 2018, 2019). Applying these measures can result in the detection of hundreds of species in a bat’s diet without losing information to host co-amplification (although it is worth noting that host co-amplification can benefit a bat dietary study by simultaneously detecting a wide range of prey taxa and confirming the predator species from faecal samples; Galan et al. 2018; Tournayre et al. 2020). Although investigations into the diets of ground-dwelling and semi-aquatic mammalian insectivores using DNA metabarcoding are less frequent, recent studies have included comparisons of primer combinations and host/diet detection (Brown et al. 2014; Esnaola et al. 2018) and those focusing on resource overlap between different insectivores (Brown et al. 2014; Biffi et al. 2017a). Studies searching for the ‘best’ primer combinations tend to have been performed on a single insectivore niche (e.g. flying or semi-aquatic). While it has been acknowledged that the best primer combination for detecting invertebrate prey in one system may not be the best for another (Tournayre et al. 2020), there has been a lack of studies investigating this directly. It is therefore important to directly compare the effect of various primers on multiple insectivores occupying different ecological systems (Corse et al. 2019).

Here we apply DNA metabarcoding to examine the diet of three mammalian insectivores with two widely used primer pairs (Zeale et al. 2011; Gillet et al. 2015) targeting the mitochondrial Cytochrome C Oxidase Subunit 1 (COI) region (chosen due to its high taxonomic coverage, resolution and well-defined reference database; Clarke et al. 2017; Elbrecht et al. 2019). These primer pairs differ in terms of prey identified (dietary constituents) and predator (host) amplification (Esnaola et al. 2018; Aldasoro et al. 2019). The three focal insectivores were chosen based on ecological niche and their proposed broad diet. The lesser horseshoe bat (*Rhinolophus hipposideros*) was used to represent a flying predator, while the pygmy shrew (*Sorex minutus*) and greater white-toothed shrew (*Crocidura russula*) were used to represent ground-dwelling predators. The diet of lesser horseshoe bats is known to be highly diverse, with 11 orders identified overall but largely dominated by Diptera and Lepidoptera as shown by both hard-part and DNA metabarcoding analyses (Aldasoro et al. 2019; Baroja et al. 2019; McAney & Fairley 1989). Their diet also changes by season and locality, demonstrating an opportunistic predatory behaviour (McAney & Fairley 1989; Baroja et al. 2019). Pygmy shrews have a diet consisting of 12 identified orders from multiple hard-part dietary analyses, with Araneae, Coleoptera and Opiliones highly represented across different parts of the species’ range (Meharg et al. 1990; Churchfield & Rychlik 2006). A recent shotgun metagenomics study (not to be confused with the metabarcoding approach used here) on five individuals also identified the importance of Lepidoptera and Acari (Ware et al. 2020). Detailed studies of the greater white-toothed shrew’s diet are limited, but Lepidoptera larvae, Araneae and Isopoda are important components of the species’ diet in Europe (Bever 1983). The species is known to catch vertebrates (including reptiles, amphibians and young small mammals; Churchfield 2008) but concrete evidence of predation is lacking. Lizards/geckos have occasionally been recovered from stomachs of the species in its African range, but it is unclear if this is due to predation or scavenging (Brahmi et al. 2012).

Focusing on these three different species, our main objective in this study was to establish whether different primer sets (or a combination of these primer sets) are appropriate for detecting different trophic niches in multiple insectivorous mammals.

## Methods

### Sample Collection and DNA Extraction

Bat faecal samples were collected non-invasively by Harrington (2018) at known bat roosts along their distribution range in the west of Ireland (Fig. S1). Sampling of bat roosts was carried out under licence from the NPWS (licence number DER/BAT 2016-29). Large sheets of plastic were laid on the ground within each roost and left for a period of one to two weeks. Droppings were collected and stored frozen at −20°C or DNA extracted within 24 hours using the Zymo Research Genomic DNA™ – Tissue MicroPrep kit following the protocol used for faecal DNA extraction in Harrington et al. (2019). Each DNA extract was identified to species level using a species-specific real-time PCR assay (Harrington et al. 2019) and identified to individual level using a panel of seven microsatellite markers originally designed by Puechmaille et al. (2005) and redesigned and optimised to work efficiently with faecal DNA by Harrington (2018) via two multiplex PCRs. Each sample was amplified, analysed and scored via three independent PCRs. A total of 24 individuals identified as *R. hipposideros* in Harrington (2018) were used in this study.

Pygmy shrews (*S. minutus*) and greater white-toothed shrews (*C. russula*) were trapped from hedgerows along secondary and tertiary roads adjacent to agricultural land in Ireland and Belle Île (France; Fig. S1). Shrews were immediately euthanised by cervical dislocation following guidelines set out by Sikes (2016) and under licences C21/2017, AE18982/I323 (Ireland) and A-75-1977 (Belle Île), and ethical approvals ST1617-55 and AREC-17-14. Carcasses were stored in separate disposable bags in a cooler until dissection later that day (max. 10 hrs). The entire gut (gastrointestinal) tract was removed and stored in absolute ethanol at a 1:4 (sample:ethanol) ratio (Egeter et al. 2015). To avoid cross-contamination, all dissections were performed on disposable bench covers and all tools were cleaned and flamed between samples. Gut contents were stored at −20°C upon returning from the field to the lab (max. 12 days). Gut tracts were defrosted on ice, removed from ethanol and air dried. Gut contents were removed from the intestines on disposable bench covers and tools were cleaned and flamed in between each sample to avoid cross-contamination. DNA was extracted from the entire gut contents using the DNeasy Power Soil Kit (Qiagen). DNA extractions were quantified using the Qubit dsDNA BR assay kit (Thermo Fisher Scientific) and subsequently diluted in molecular grade water to 10–15 ng/μl. A subset of 12 *C. russula* (10 from Ireland, and 2 from Belle Île) and 15 *S. minutus* (10 from Ireland, and 5 from Belle Île) samples were chosen for this study. In total, 51 insectivores were analysed, including 27 ground-dwelling and 24 flying individuals. Details of the samples used can be found in Table S1.

### Polymerase Chain Reaction (PCR)

DNA extracts were amplified using two primer sets targeting different short fragments of the mtDNA COI gene. The Zeale primers (ZBJ-ArtF1c 5’-AGATATTGGAACWTTATATTTTATTTTTGG-3’ and ZBJ-ArtR2c 5’-WACTAATCAATTWCCAAATCCTCC-3’ Zeale et al. 2011) were used to amplify a 157bp section of COI, and the Gillet primers ((modified LepF1 (Hebert et al. 2003) 5’-ATTCHACDAAYCAYAARGAYATYGG-3’)) and ((EPT-long-univR (Hajibabaei et al. 2011) 5’-ACTATAAAARAAAATYTDAYAAADGCRTG-3’)) were used to amplify 133bp of COI. The two pairs of primers will be referred to as the Zeale and Gillet primer sets and datasets from here on. A set of 24 unique eight base pair multiplex identifiers (MID) tags were added to the Zeale and Gillet primer sets to allow for the multiplexing of samples into a single sequencing run. A different set of 24 unique MID tags were used for each primer pair.

The PCR mix for both Gillet and Zeale primer sets contained 12.5 μl Qiagen Multiplex PCR Mastermix, 1 μl of each primer (5 μm), 7.5 μl of molecular grade water and 3 μl of DNA template (molecular grade water for negative controls). PCR conditions for the Zeale primers included an initial denaturation at 95°C for 15 minutes, followed by 40 cycles of 95°C for 20 seconds, 55°C for 30 seconds and 72°C for one minute, followed by a final extension at 72°C for seven minutes (Aizpurua et al. 2018; Alberdi et al. 2018). PCR conditions for Gillet primers were trialled from Esnaola et al. (2018) but amplified a non-target region of DNA approximately 200 bp and 500 bp larger than the target region in *S. minutus* samples. The PCR conditions were altered to a two-stage PCR with higher annealing temperatures to increase specificity and decrease amplification of non-target fragments. The altered PCR conditions for Gillet primers involved an initial denaturation at 95°C for 15 minutes followed by 10 cycles of 94°C for 30 seconds, 49°C for 45 seconds and 72°C for 30 seconds, followed by 30 cycles of 95°C for 30 seconds, 47°C for 45 seconds, 72°C for 30 seconds followed by a final extension of 72°C for 10 minutes. The PCRs were run in triplicate, subsequently pooled and the success of the reactions was determined by electrophoresis on a 1.2% agarose gel, which included the two negative control PCR products.

Library preparation, sequencing and bioinformatic steps are provided in Appendix 1 of the Supplementary Material.

### Taxonomic Identification and Range

The number of MOTUs identified and taxonomically assigned to different levels were compared between datasets using sequence clustering thresholds 95% and 98% to determine the capabilities of both primers and the overall effect of the clustering threshold. The final clustering thresholds were chosen based on the number and proportion of MOTUs that were taxonomically assigned. The clustering threshold chosen was the value with the highest proportion of MOTUs assigned to species and genus level, with reduced proportions of MOTUs restricted to order and family. In addition to this, the clustering values commonly used in the literature were also taken into account for our choice (Alberdi et al. 2018).

The taxonomic range was compared at each taxonomic level between primer sets and considered separately for both bats and shrews to establish if one primer was suited to a particular predator diet. To assess the ability of each primer to detect unique taxa, the overlap of accurately identified taxa was measured between Zeale and Gillet primers for bats and shrews at order, family, genus and species level.

### Alpha Diversity

The samples represented by the combined effort of both Zeale and Gillet here have an extra advantage of increased sequencing depth. To account for this in alpha diversity measures, samples (and groups of samples) were rarefied to an equal sequencing depth to achieve a more accurate comparison. Samples were rarefied to the lowest sampling depth (1110 reads) before alpha diversity measures (species richness and Shannon diversity) were calculated. To account for any stochastic results from rarefying samples, this process was repeated 100 times and the average alpha diversity scores were taken for each metric. Significant differences in alpha diversity between groups were identified using ANOVA and a Tukey post-hoc test.

The samples were then merged according to mammal species and primer used by summing the reads for each MOTU. The merged samples were then rarefied to the lowest read depth of said merged samples (105501 reads). The niche width of each mammal species amplified by different primers was measured using the standardised Levin’s index, Shannon diversity index (for details on measurements see Razgour et al. 2011) and Pielou’s eveness index using the R packages vegan and spaa (Zhang 2016).

### Beta Diversity

Data were normalised by transforming sequence counts into relative read abundances per sample and a distance matrix was created for the dataset using the Bray-Curtis dissimilarity method. Data were visualised using a Non-metric multidimensional scaling (NMDS) ordination plot. To determine any compositional difference in prey taxa identified between consumer species and/or primer used, permutational multivariate analysis of variance (PERMANOVA) were performed with 10000 permutations using the *adonis2* function in the *vegan* package in R. To be certain that any composition differences were not due to differences between homogeneity of dispersion within groups, the multivariate distances of samples to the group centroid was measured using the *betadisper()* function. All beta diversity estimates described here were repeated with MOTUs agglomerated to species, genus, family, and order levels.

### Hierarchical Clustering

Hierarchical clustering was performed to show how the chosen primer affects the grouping of samples. Clustering was performed on each sample using the *hclust()* function in R, with the UPGMA method. Clustering was also performed on samples grouped according to predator species and primer using the average Relative Read Abundance (RRA) values.

### Random Forest Classifier

While different primers will amplify different taxonomic groups, it is desirable to determine which of the tested primers will amplify a greater range of taxa important to characterising the diet of that predator species. The random forest classification (RFC) is a supervised learning method that classifies samples (such as prey composition) to their source, estimates the level of importance of each prey item to that classification and determines the accuracy of that classification (Breiman et al. 2001). Here, RFC models were run to firstly determine which primer amplifies taxa that are most appropriate for classifying samples to predator species, and then again secondly to classify samples to the correct predator species based on the prey composition.

RFCs were performed on samples using the *randomForest* R package (Liaw and Weiner 2002) using 10,000 trees. The out-of-bag (OOB) error was used to measure the accuracy of classification of samples to their correct group. The most important prey taxa contributing to classification of samples were established using the ‘Mean Decrease Mini’ values.

## Results

### Bioinformatics and MOTU filtering

The MiSeq sequencing run produced 18,527,116 sequence reads; 48.4% associated with bat samples and 49.5% associated with shrew samples. A sequence clustering threshold of 98% was used for downstream analyses. This clustering threshold identified MOTUs that had the highest species and genus level assignment rates, with lower levels of assignment restricted to family and order level (Fig. S2). This threshold has been used by many other studies using the COI region for invertebrate detection (Alberdi et al. 2018).

The dataset utilising the sequence clustering threshold at 98% similarity yielded 9,647 non-singleton MOTUs and 7,698 non-singleton MOTUs for the Gillet and Zeale datasets, respectively. In the negative controls, the Gillet dataset returned 5,085 reads from the *Chiroptera* order (<0.13% of all *Chiroptera* reads) and 56 reads from *Homo sapiens* (~3.25% of all human reads). The *Rhinolophidae* reads in the negative control accounted for only 0.08% of all host reads across the entire dataset amplified by the Gillet primer set. These MOTUs were excluded from further analyses.

After removing MOTUs according to filtering criteria and samples with low read counts, the Gillet dataset contained 945 MOTUs across 22 *R. hipposideros*, 7 *C. russula* and 15 *S. minutus*, with an average read depth of 37,555 reads per individual. The Zeale dataset contained 929 MOTUs across 23 *R. hipposideros*, 4 *C. russula* and 11 *S. minutus* with an average read depth of 159,589 per individual. Rarefaction curves showed that all prey taxa were detected between 1000 and 5000 reads for each sample (Fig. 1A: inset) and the depth_cov(., qvalue = 1) function showed a sample coverage of >97% for Zeale and >98% for Gillet.

**Figure 1.**
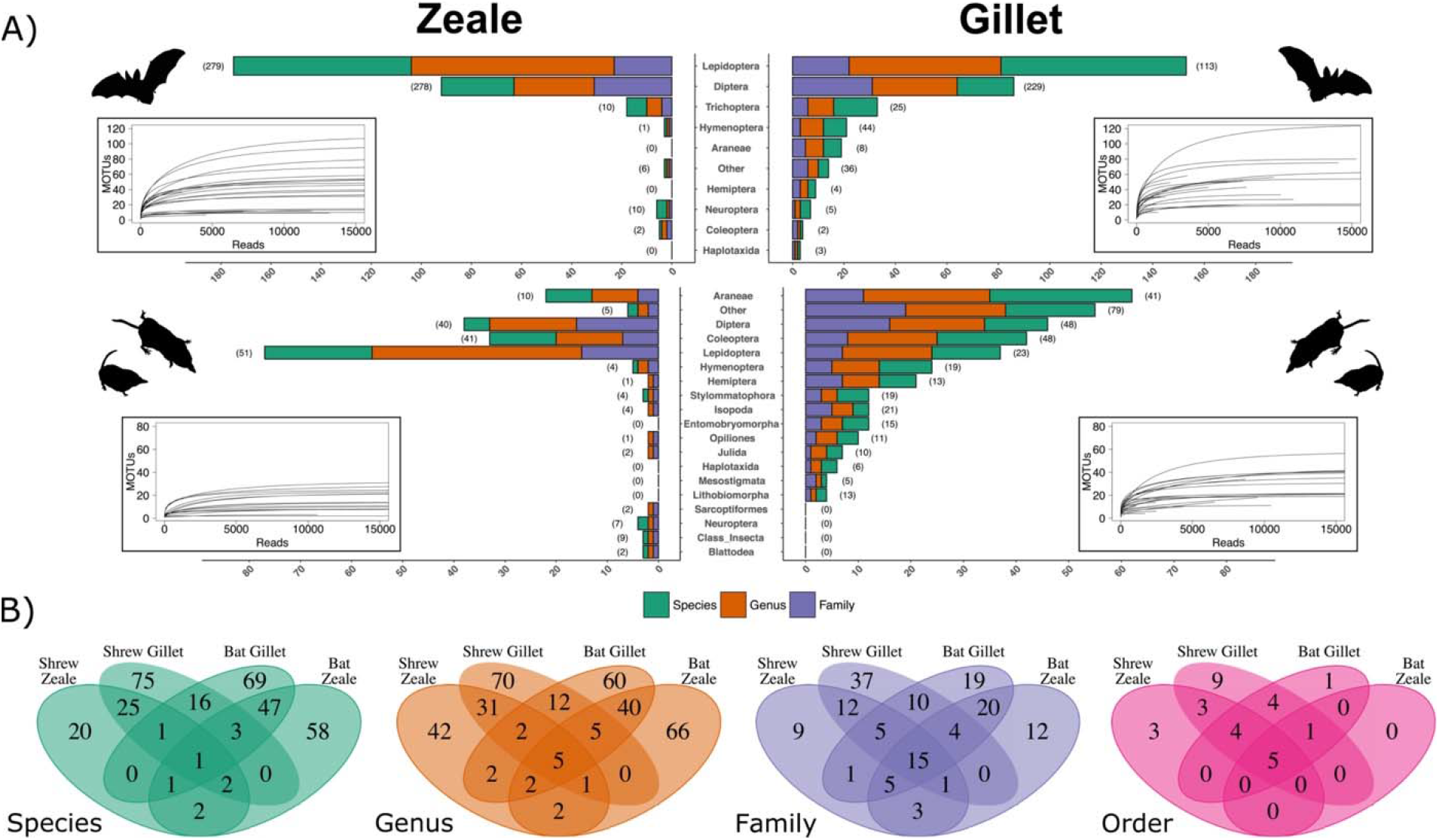
Prey detection of Zeale and Gillet primers in bats (*R. hipposideros*) and shrews (*C. russula* and *S. minutus*). A) Bar plots showing the number of prey species, genera and families detected in each of the most abundant prey orders. Numbers in parentheses represent the number of MOTUs detected in each order. Inset plots are rarefaction curves estimating that 1000 to 5000 reads are required to capture total species richness per sample. B) Venn diagrams showing how many of the detected prey taxa are shared between the Zeale and Gillet primers.

### Taxonomic Identification and Range

Both primers detected similar numbers of MOTUs; the Gillet primers detected MOTUs that were taxonomically assigned to 240 species, 230 genera, 129 families and 27 orders. The Zeale primers detected MOTUs that were taxonomically assigned to 160 species, 198 genera, 87 families and 16 orders.

Both primers detected a similar number of prey taxa in bats (**Fig. 1**). The majority of taxa detected belong to the orders Lepidoptera and Diptera, with some taxa within the Trichoptera order. Gillet also detected a small number of taxa from Hymenoptera and Araneae in the bat diet. Haplotaxida were detected by the Gillet primers, but this is likely due to environmental contamination. Although both primers detected the majority of species within Lepidoptera and Diptera in bat samples, there was a relatively even distribution of taxa detected by one and both primers (**Fig. 1B**).

There was a more prominent difference between primers for taxa detection in shrews. The majority of taxa identified by Zeale were within the orders Lepidoptera, Diptera, Coleoptera and Araneae (**Fig. 1A**). Gillet detected taxa from a much wider order of terrestrial invertebrates (such as Haplotaxida, Hemiptera, Stylomatophora, Isopoda and more) that are considered important in the diet of shrews (Pernetta 1976; Churchfield and Rychlik 2006). Additionally, Gillet detected substantially more species, genera, families and orders that Zeale could not (**Fig. 1B**). The three orders detected by only Zeale are Sacoptiformes, Neuroptera and Blattodea which contained only 2, 7 and 2 MOTUs respectively.

As expected, the only primer set here to detect vertebrate DNA was the Gillet primers. Between 89% and 99% of reads in bats were of vertebrate origin and between 0.81% and 99% of reads in shrew samples were of vertebrate origin.

### Composition of Diet

The average relative read abundance (RRA) of prey order in *R. hipposideros* diet did not dramatically change between primer sets (**Fig. 2B**). Both primers showed that the diet mostly consisted of Diptera and Lepidoptera, but only the Gillet primers showed a noticeable proportion of the diet consisting of Hymenoptera and Trichoptera. Using a combination of both primers showed a stronger similarity than using Zeale primers alone, complementing the hierarchical clustering (**Fig. 2A**).

**Figure 2.**
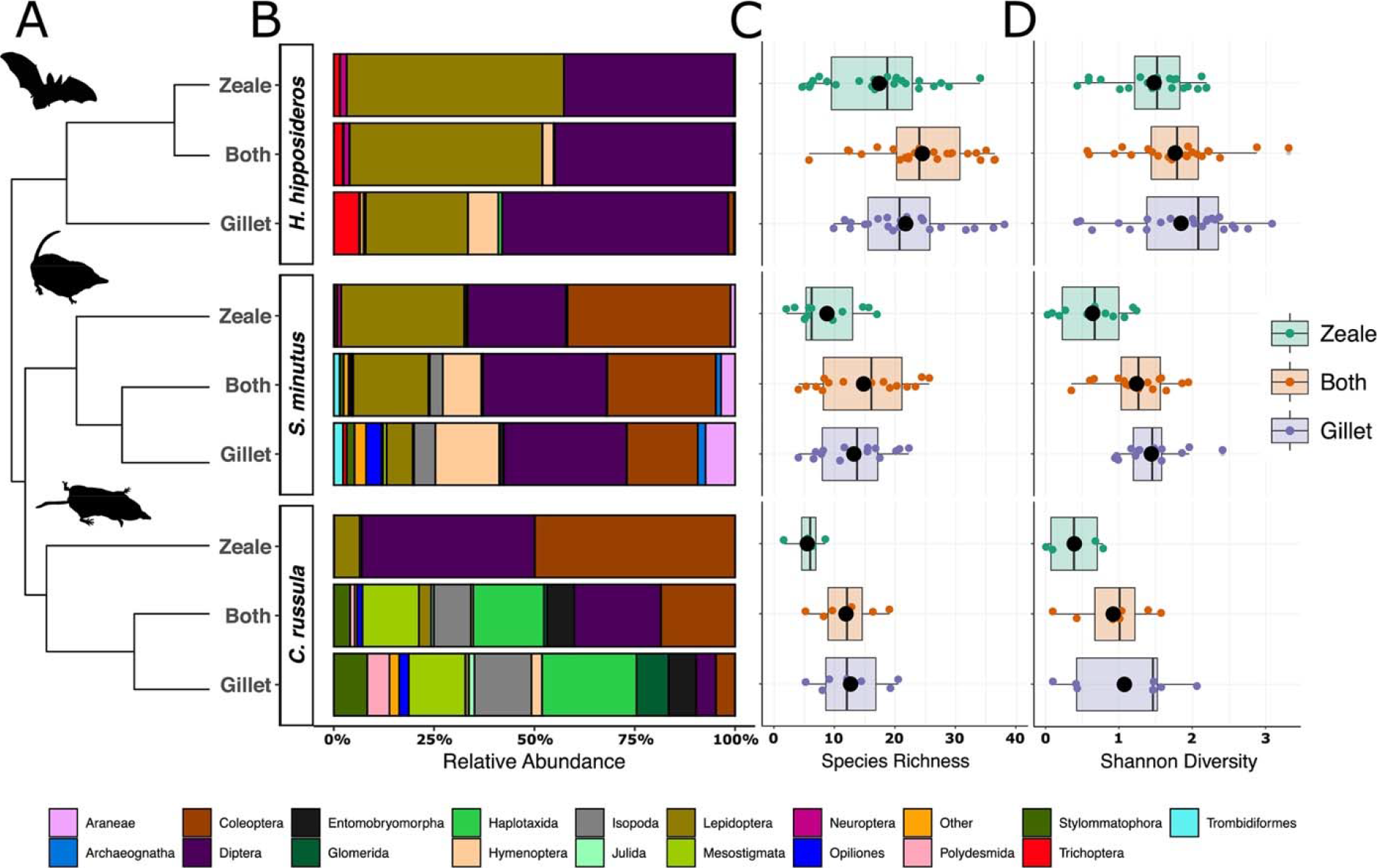
A) Hierarchical clustering of mammal species amplified with Zeale, Gillet and Both (Zeale and Gillet combined) primer sets. B) Average relative abundance of prey orders. C) Invertebrate species richness recovered for each analysed mammal species D) Shannon diversity per mammal species.

When used individually, the Zeale and Gillet primer sets suggested that a large proportion of the diet of *S. minutus* consisted of Lepidoptera, Diptera and Coleoptera (**Fig. 2B**). Only the Gillet primers suggested the additional importance of other orders such as Araneae, Hymenoptera, Isopoda, Opilliones and Trombidiformes as contributing to the diet of *S. minutus*. Using both primers to determine the diet of *S. minutus* demonstrated a strong influence by Gillet, complementing the hierarchical clustering (**Fig. 2A**), but with larger proportions of Lepidoptera, Diptera and Coleoptera.

*Crocidura russula* showed the largest differences in diet when analysed by Zeale or Gillet primers (**Fig. 2B**). Again, Zeale was restricted to Lepidoptera, Diptera and Coleoptera. Gillet suggested the importance of terrestrial invertebrates such as Haplotaxida, Glomerida, Isopoda, Mesostigmata and Stylomatophora. Using a combination of both primers resembled the diet suggested by Gillet alone, complementing the hierarchical clustering (**Fig. 2A**).

### Alpha Diversity

After agglomerating taxa to their highest taxonomic level, the Gillet and Zeale datasets consisted of 425 and 371 prey MOTUs respectively, with a combined richness of 660 MOTUs (**Table 1**). The mean alpha diversity measures were higher in *R. hipposideros* compared to shrews, with *S. minutus* marginally higher than *C. russula* (**Figs 2C and 2D**). For species richness, Tukey post-hoc comparison of means showed that *R. hipposideros* samples amplified with both primers had an average of between 13.36 and 20.5 more MOTUs detected than all *C. russula* samples (all adjusted p-values < 0.01), between 11.2 and 17.3 MOTUs more than all *S. minutus* samples (all adjusted p-values < 0.01) and 8.6 more MOTUs than *R. hipposideros* amplified with Zeale (adjusted p-value <0.02). *Rhinolophus hipposideros* samples amplified with Gillet primers had an average of 14.2 and 17.5 more MOTUs than *S. minutus* and *C. russula* samples amplified with Zeale, respectively (adjusted p-value <0.001).

**Table 1.**
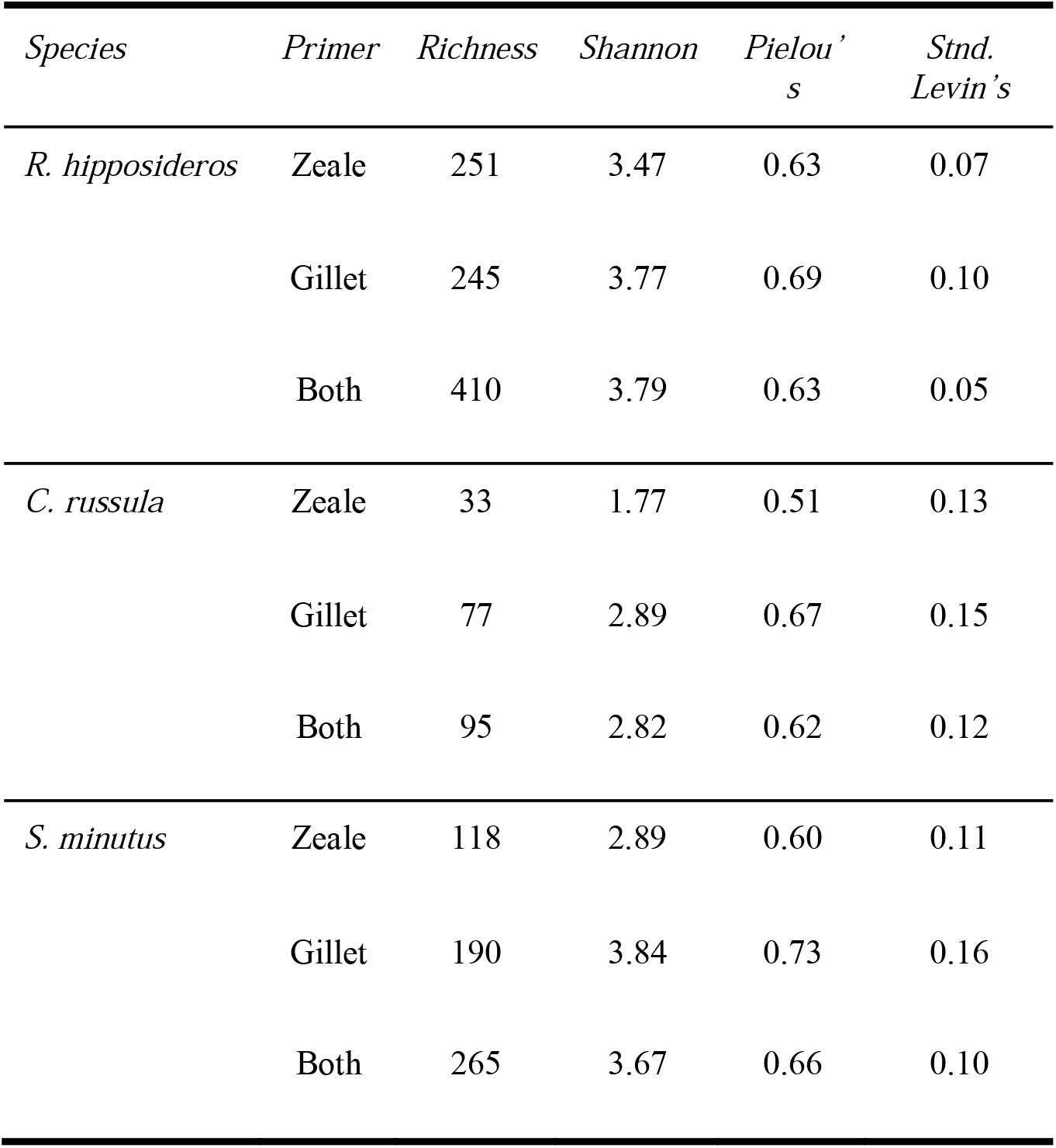
Alpha Diversity Measures for each primer set (Zeale and Gillet) and Both (Zeale and Gillet combined). Pielou’s is a measure of evenness. Standardised Levin’s is typically used as a measure of niche breadth.

For Shannon diversity, the Tukey post-hoc comparison of means showed significantly lower diversity (adjusted p-value < 0.05) in *C. russula* amplified by Zeale primers compared to all bat samples, and *S. minutus* samples amplified with Gillet primers. Amplifying *C. russula* samples with both primers produced significantly lower diversity values than *R. hipposideros* amplified with either Gillet or both primers. *Sorex minutus* samples amplified with Zeale primers had significantly lower values than all *R. hipposideros samples.* One notable difference is the significantly lower Shannon diversity in *S. minutus* samples amplified with Zeale compared to Gillet (adjusted p-value = 0.19).

### Beta Diversity

PERMANOVAs estimated a significant, but minor, difference in the composition of prey detected in *R. hipposideros* when using Gillet vs Zeale (R^2^ = 0.08, Pr(>F) = 0.001) and Gillet vs Both (R^2^ = 0.05, Pr(>F) = 0.001) but not for Zeale vs Both (R^2^ = 0.006, Pr(>F) = 1). The NMDS plot (**Fig. 3**) showed that bats amplified with Gillet, Zeale and both primers clustered close together which also suggested that compositional differences are likely minor. There was also a minor, but significant, difference in the prey composition detected in shrews when comparing Gillet vs Zeale samples (R^2^ = 0.038, Pr(>F) = 0.029) (also seen in **Fig. 3**). Each primer set could detect a composition difference between *R. hipposideros* and shrews (R^2^ = 0.044 – 0.067, Pr(>F) < 0.01), which is a visibly clear pattern in the NMDS plot in **Fig. 3**.

**Figure 3.**
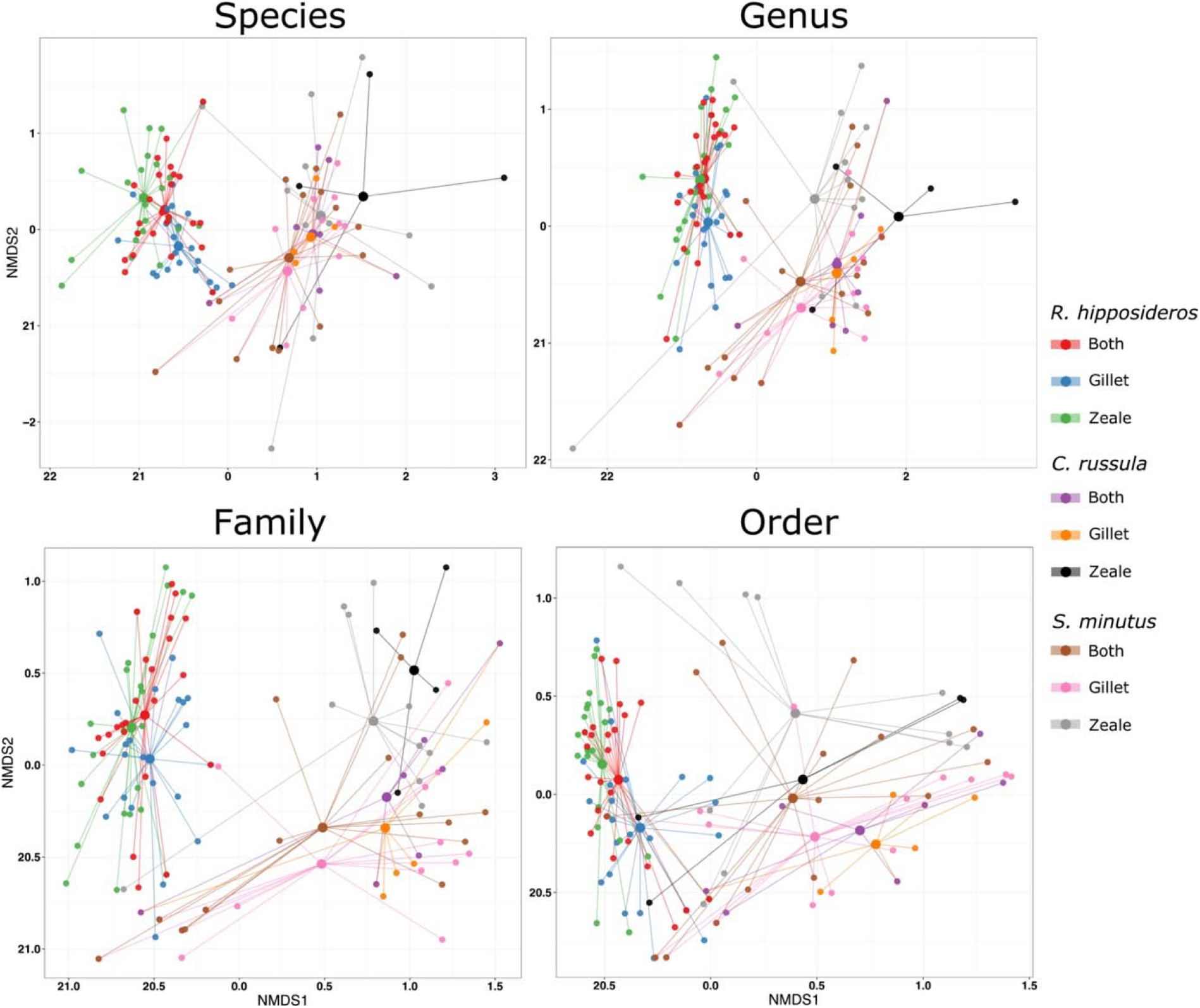
NMDS plots of samples when MOTUs are agglomerated according to species, genus, family and order.

The Tukey pairwise comparison showed no difference in the homogeneity of these tested groups, but the permutest showed a difference between *S. minutus* amplified with Zeale primers against all *C. russula* samples, which may have influenced the PERMANOVA results. The permutest also showed a difference between the homogeneity of *C. russula* amplified with Zeale compared to either Gillet (p < 0.01) or Both primers (p < 0.001). These differences should be considered while interpreting compositional differences as homogeneity can influence PERMANOVA results.

*R. hipposideros* mainly predates on Diptera and Lepidoptera (**Fig. 2B**), which may explain why they remain a tight cluster in the NMDS plots as MOTUs are agglomerated up to order level (**Fig. 3**). Although shrews (particularly *S. minutus*) also predate on Diptera and Lepidoptera (**Fig. 2B**), they remain distinct from *R. hipposideros* when MOTUs were agglomerated to species level. As MOTUs are agglomerated to higher levels, the coordinates of some shrews migrate and cluster closer to *R. hipposideros.* This suggests that there are common prey orders between the three insectivore species, but bats and shrews still predate on different species, genera and families within these common prey orders.

### Random Forest Classifier

RFC models were able to classify samples as originating from *R. hipposideros* or shrews with an accuracy of 100% using Zeale, 88.64% using Gillet, and 93.48% using both. Amongst the top 20 most important taxa (MOTUs with the highest Mean Decrease Mini values) for classifying samples to bat or shrew, the most common prey order was Diptera and Lepidoptera for each primer used.

The accuracy was much lower for classifying samples to *C. russula* or *S. minutus* using Zeale (73.33%), Gillet (68.18%) or both (68.18%). The top 20 taxa for classifying species of shrew mainly consisted of taxa within Lepidoptera and Coleoptera when amplified using Zeale primers. Using Gillet, or both primers, the top 20 taxa were distributed more evenly amongst more orders such as Haplotaxida, Opiliones, Stylommatophora and Diptera.

Bat samples could be classified to Zeale or Gillet with a high accuracy of 93.33%, while the accuracy to classify between Gillet and both primers decreased to 73.91%, and between Zeale and both decreased to 70.21%. Shrew samples could be classified between Zeale and Gillet with a lower accuracy of 83.78%. However, accuracy drastically decreased when classifying shrews between Zeale and Both primers (54.05%) or between Gillet and both primers (2.27%). Full details on the 20 taxa with the highest mean Decrease Gini values can be found in Tables S2–S13.

## Discussion

Here we show that two different COI primer sets performed differently for detecting invertebrate prey composition across a broad ecological range, meaning that primer choice will have a significant impact on ecological inferences from the data generated with them. Primer comparisons for determining the diet in insectivorous mammals have previously been performed on single species or multiple species within the same ecological niche (e.g. bats; Tournayre et al. 2020). Here, we compared two widely used primer sets (Zeale and Gillet) on multiple mammals occupying different niches and demonstrated that while one primer set captured the breadth of prey for ground-dwelling shrews, both primer sets were required to fully capture the diet of bats within the studied systems.

When comparing the Zeale and Gillet primer sets, the first obvious and major advantage of the Zeale primers was that there was practically no host amplification, meaning that all information retained by the Zeale primer pair represents potential prey. In contrast, the Gillet primers co-amplified large amounts of host DNA (up to 99% in some samples), which has also been observed in previous studies (Baroja et al. 2019; Esnaola et al. 2018; Galan et al. 2018). The varied amount of host amplification between samples in this study highlights that rates of host amplification may be unpredictable to an extent. Host amplification affected *S. minutus* less than *R. hipposideros* and *C. russula*, and some technical and biological issues should be taken into account when analysing the difference found between species in regard to host amplification. For example, considering that the shrew samples were gut contents from dissection, ‘empty’ stomachs may have influenced the higher rate of host DNA amplification in the absence of prey DNA in some predators.

Apart from host and human DNA, the Gillet primers detected trace amounts of DNA from other vertebrates such as bank voles (*Myodes glareolus*), cattle (*Bos taurus*) and pig (*Sus scrofa*). These taxa contributed to between 2 and 16 reads in total, likely through secondary detection from invertebrate prey coming into contact with other vertebrates or their excrement before consumption. This is an unsurprising result as previous studies have detected various species of birds, mammals and amphibians with the Gillet primers (Biffi et al. 2017b; Esnaola et al. 2018; Galan et al. 2018). Host amplification is not desirable here, but the capability to amplify vertebrate DNA is beneficial to determine if the invasive *C. russula* (in Ireland) are consuming local vertebrate taxa (McDevitt et al. 2014).

This level of host amplification means that the average number of reads attributed to invertebrates in each sample was approximately three times lower in Gillet compared to Zeale. An insufficient read depth will reduce the likelihood of detecting the entire prey community, but rarefaction estimates suggested that the majority of prey were detected with a sequencing depth of between 1,000 and 5,000 reads (**Fig. 1A**). Despite the reduced read depth for prey using Gillet, more samples satisfied the filtering criteria when amplified with Gillet rather than Zeale. This is due to the Gillet primers ability to amplify a wider range of taxa, including an additional 14 orders (**Fig. 2B**). Many of these additional orders constitute a large portion of different shrew species’ diet, such as slugs/snails (Stylommatophora), spiders (Araneae), woodlice (Isopoda), millipedes (Polydesmida) and worms (Haplotaxida) (**Fig. 2B**; Pernetta, 1976). These results showed that after removing host sequences, Gillet primers provided more information on invertebrate prey than Zeale without using blocking primers once sufficient sequencing depth is achieved. Furthermore, blocking primers can mitigate host DNA amplification but requires more time to design and test as they might also block amplification of target prey taxa (Piñol et al. 2015) and would be particularly challenging when investigating multiple species simultaneously as undertaken here.

The Zeale primers are extensively used and have proved very efficient in determining the diet of bats (Vesterinen et al. 2018), but this trial showed that in terrestrial insectivores they are still mostly limited to the three orders: Coleoptera, Diptera and Lepidoptera. They are more suitable for bats, as even the Gillet primers with their wider taxonomic range show that Diptera and Lepidoptera are the main constituents of their diet (**Fig. 2B**) and is in agreement with previous studies on *R. hipposideros* (Aldasoro et al. 2019; Baroja et al. 2019). Due to Zeale’s high affinity to Coleoptera, Diptera and Lepidoptera, shrew diets were biased towards these orders (**Fig. 2B**). In addition, the rate of shrew samples filtered out due to low read counts was much higher than with Gillet. It was evident from this study (and previous studies; Ware et al. 2020) that shrews also rely on other terrestrial invertebrate orders such as Gastropoda, Isopoda and Haplotaxida (**Fig. 2B**). Zeale’s inability to detect these taxa means that many shrew samples were filtered out during bioinformatic processing. Using the Gillet primers, some of the orders listed as substantial in the diet of shrews were also detected in the bat diet (i.e. Araneae and Haplotaxida) (**Fig. 2B**). While Aranea have previously been identified in the diet of bats (McAney and Fairley 1989), Haplotaxida have not. This unexpected detection is likely a result of environmental contamination (Aldasoro et al. 2019). In each of the *R. hipposideros* roosts sampled in Ireland, large sheets of plastic were laid down to collect faecal samples and left exposed for a period of up to two weeks. Therefore, organisms coming into contact with the samples from nearby guano piles during this time may explain their detections, as Haplotaxida have been reported in bat guano elsewhere (Novak et al. 2014).

Recent studies suggest that using more than one primer will cover a wider range of taxa and give a more informative overview of the diet of these animals (Esnaola et al. 2018). This is true considering they both amplify unique taxa. For example, even though Zeale and Gillet both amplify MOTUs within the orders Diptera and Lepidoptera, they each amplify several unique MOTUs/species within each (Baroja et al. 2019). In addition, the RFC analysis showed that there is still a relatively high accuracy differentiating bat samples amplified with Zeale or Gillet (>90%) and only decreased in accuracy to ~70% when including samples amplified with both primers. This supports both that primers are contributing relatively evenly in detecting unique components of the diet of bats. The composition of the detected diet of shrews using both primers appeared heavily influenced by the Gillet set, rather than Zeale (**Figs 2 and 3**), which was particularly apparent at the order level. The RFC analysis had a very low efficiency differentiating samples that had been amplified with Gillet primers or both (2.27%). In addition, when considering the same finite number of sequences that can be generated, combining the Zeale and Gillet data increased diversity of shrew prey detected compared to Zeale alone but did not significantly increase diversity compared to using Gillet alone (**Fig. 2C**). This was likely due to Gillet detecting more substantial components of a shrew’s diet such as slugs/snails, spiders, woodlice, millipedes and worms. A combined effect of primers will also restrict dietary studies to frequency/occurrence-based analyses. Although many studies stick to a more conservative frequency based interpretations of dietary data, relative read abundance (RRA) can still accurately represent the proportions of prey in an animal’s diet at the population level (Deagle et al. 2019). Combining both primers used here (and in future studies) will require the sequencing depth to be normalised between the primer datasets if RRA methods are to be used since the proportions of prey taxa become skewed in favour of Coleoptera, Diptera and Lepidoptera (Error! Reference source not found.**B**).

Including both primers in a full-scale analysis will obviously increase costs and labour so the research question to be addressed becomes the critical component when deciding which primer set(s) combination to use when investigating mammalian insectivore diet. For the species considered here, the Gillet primers amplify a wider range of taxa and may be sufficient to address ecological questions around dietary composition (e.g., spatial and temporal shifts) and competition/overlap between species (particularly for shrews). However, given the importance of bats in providing ecosystem services, and their potential role as ‘natural samplers’ (Siegenthaler et al. 2019) for undertaking invertebrate surveying, multiple primer sets would be required, particularly when individual pest species may need to be identified and/or monitored (Baroja et al. 2019).

## Authors’ contributions

ADM, SSB, TGC and DBO’M conceived and designed the study. Bat faecal sampling was part of APH, D’ON and DBO’M’s project on non-invasive genetic monitoring of lesser horseshoe bats. Shrew sampling was part of SSB, REA and ADM’s project on dietary and microbial associations between shrew species in Ireland. SSB, TGC and NGS performed the laboratory work and bioinformatics associated with the DNA metabarcoding. SSB and TGC analysed the data. SSB, TGC and ADM wrote the paper, with all authors contributing to editing, discussions and approval of the final manuscript.

## Acknowledgements

SSB was supported by a Pathway to Excellence PhD Scholarship from the University of Salford and TGC was supported by a Waterford Institute of Technology and Environmental Protection Agency (EPA) cofund PhD Scholarship, funded under the EPA Research Programme 2014-2020. The EPA Research Programme is a Government of Ireland initiative funded by the Department of the Environment, Climate and Communications. It is administered by the EPA, which has the statutory function of co-ordinating and promoting environmental research. Thank you to the local National Park and Wildlife Service (NPWS) conservation rangers for sampling bat droppings. Fieldwork in relation to shrew sampling was supported by grants from the Vincent Wildlife Trust and The Genetics Society awarded to SSB and ADM. Laboratory work was supported by grants from Bat Conservation Ireland and a University of Salford Internal Research Award awarded to DBO’M and ADM, and a University of Salford Pump Priming award to ADM. NGS also thanks FCT/MCTES for the financial support to CESAM (UID/AMB/50017/2019), through national funds.

## Data Accessibility

All bioinformatic steps and scripts can be found on github (https://github.com/ShrewlockHolmes). Raw sequence data will be made publicly available upon publication.

## References

Aizpurua O, Budinski I, Georgiakakis P, Gopalakrishnan S, Ibañez C, Mata V, Rebelo H, Russo D, Szodoray-Parádi F, Zhelyazkova V, Zrncic V, Gilbert MTP, Alberdi A (2018) Agriculture shapes the tropic niche of bats preying on multiple pest arthropods across Europe: Evidence from DNA metabarcoding. Molecular Ecology 27(3):815–825. doi: 10.1111/mec.14474.

Alberdi A, Aizpurua O, Gilbert MTP, Bohmann K (2018) Scrutinizing key steps for reliable metabarcoding of environmental samples. Methods in Ecology and Evolution 9(1):134–147. doi: 10.1111/2041-210X.12849.

Alberdi A, Aizpurua O, Bohmann K, Gopalakrishnan S, Lynggaard C, Nielsen M, Gilbert MTP (2019) Promises and pitfalls of using high ◻ throughput sequencing for diet analysis. Molecular Ecology Resources 19(2):327–348. doi: 10.1111/1755-0998.12960.

Aldasoro M, Garin I, Vallejo N, Baroja U, Arrizabalaga-Escudero A, Goiti U, Aihartza J (2019) Gaining ecological insight on dietary allocation among horseshoe bats through molecular primer combination. Plos One 14(7):e0220081. doi: 10.5061/dryad.hm3v1dm.

Baroja U, Garin I, Aihartza J, Arrizabalaha-Escudero A, Vallejo N, Aldasoro M, Goiti U (2019) Pest consumption in a vineyard system by the lesser horseshoe bat (*Rhinolophus hipposideros*). Plos One 14 (7):e0219265.

Bever K (1983) Zur Nahrung der Hausspitzmaus, *Crocidura russula* (Hermann 1780). Saugetier Mittl 31:13–26

Biffi M, Laffaille P, Jabiol J, André A, Gillet F, Lamothe S, Michaux JR, Buisson L (2017a) Comparison of diet and prey selectivity of the Pyrenean desman and the Eurasian water shrew using next-generation sequencing methods. Mammalian Biology 87:176–184. doi: 10.1016/j.mambio.2017.09.001.

Biffi M, Gillet F, Laffaille P, Colas F, Aulagnier S, Blanc F, Galan M, Tiouchichine M-L, Némoz M, Buisson L, Michaux JR (2017b) Novel insights into the diet of the Pyrenean desman (*Galemys pyrenaicus*) using next-generation sequencing molecular analyses. Journal of Mammalogy 98(5):1497–1507. doi: 10.1093/jmammal/gyx070.

Brahmi K, Aulagnier S, Slimani S, Mann CS, Doumandji S, Baziz B (2012) Diet of the Greater white-toothed shrew *Crocidura russula* (Mammalia: Soricidae) in Grande Kabylie (Algeria). Italian Journal of Zoology 79(2):239–245. doi: 10.1080/11250003.2011.625449.

Breiman L (2001) Random forests. Machine Learning 45(1):5–32.

Browett SS, O’Meara DB, McDevitt AD (2020) Genetic tools in the management of invasive mammals: recent trends and future perspectives. Mammal Review 50(2):200–210. doi: 10.1111/mam.12189.

Brown DS, Burger R, Cole N, Vencatasamy D, Clare E, Montazam A, Symondson WOC (2014) Dietary competition between the alien Asian Musk Shrew (*Suncus murinus*) and a re-introduced population of Telfair’s Skink (*Leiolopisma telfairii*). Molecular Ecology 23(15):3695–3705. doi: 10.1111/mec.12445.

Churchfield S (2008) Greater white-toothed shrew, in Harris, S. and Yalden, D. W. (eds) Mammals of the British Isles handbook. 4th edn. The Mammal Society, pp. 280–283.

Churchfield S and Rychlik L (2006) Diets and coexistence in *Neomys* and *Sorex* shrews in Białowieża forest, eastern Poland. Journal of Zoology 269(3):381–390. doi: 10.1111/j.1469-7998.2006.00115.x.

Clarke LJ, Beard JM, Swadling KM, Deagle BE (2017) Effect of marker choice and thermal cycling protocol on zooplankton DNA metabarcoding studies. Ecology and Evolution 7(3):873–883. doi: 10.1002/ece3.2667.

Corse E, Tougard C, Archambaud-Suard G, Agnèse JF, Messu Mandeng FD, Bilong Bilong CF, Duneau D, Zinger L, Chappaz R, Xu CCY, Meglécz E, Dubut V (2019) One◻locus◻several◻primers: A strategy to improve the taxonomic and haplotypic coverage in diet metabarcoding studies. Ecology and Evolution 9:4603–4620. doi: 10.1002/ece3.5063.

Deagle BE, Thomas AC, McInnes JC, Clarke LJ, Vesterinen EJ, Clare EL, Kartzinel TR, Eveson JP (2019) Counting with DNA in metabarcoding studies: How should we convert sequence reads to dietary data?. Molecular Ecology 28(2):391–406. doi: 10.1111/mec.14734.

Egeter B, Bishop PJ, Robertson BC (2015) Detecting frogs as prey in the diets of introduced mammals: a comparison between morphological and DNA-based diet analyses. Molecular Ecology Resources 15(2):306–316. doi: 10.1111/1755-0998.12309.

Elbrecht V, Braukmann TWA, Ivanova NV, Prosser SWJ, Hajibabaei M, Wright M, Zakharov EV, Hebert PDN, Steinke D (2019) Validation of COI metabarcoding primers for terrestrial arthropods. PeerJ 7:e7745. doi: 10.7717/peerj.7745.

Esnaola A, Arrizabalaga-Escudero A, González-Esteban J, Elosegi A, Aihartza J (2018) Determining diet from faeces: Selection of metabarcoding primers for the insectivore Pyrenean desman (*Galemys pyrenaicus*). PLoS ONE 13 (12):e0208986. doi: 10.5061/dryad.bt211nm.Funding.

Galan M, Pons J-B, Tournayre O, Pierre É, Leuchtmann M, Pontier D, Charbonnel N (2018) Metabarcoding for the parallel identification of several hundred predators and their prey◻: Application to bat species diet analysis. Molecular Ecology Resources 18:474–489. doi: 10.1111/1755-0998.12749.

Gillet F, Tiouchichine M-L, Galan M, Blanc F, Némoz M, Aulagnier S, Michaux JR (2015) A new method to identify the endangered Pyrenean desman (*Galemys pyrenaicus*) and to study its diet, using next generation sequencing from faeces. Mammalian Biology 80(6):505–509. doi: 10.1016/j.mambio.2015.08.002.

Hajibabaei M, Shokralla S, Zhou Z, Singer GAC, Baird DJ (2011) Environmental barcoding: A next-generation sequencing approach for biomonitoring applications using river benthos. PLoS ONE 6(4). doi: 10.1371/journal.pone.0017497.

Harrington AP (2018) The Development of Non-Invasive Genetic Methods for Bats of the British Isles. Unpublished PhD thesis, Waterford Institute of Technology.

Harrington AP, O’Meara DB, Aughney T, McAney K, Schofield H, Collins A, Deenen H, O’Reilly C (2019) Novel real-time PCR species identifcation assays for British and Irish bats and their application to a non-invasive survey of bat roosts in Ireland. Mammalian Biology 99:109–118. doi: 10.1016/j.mambio.2019.10.005.

Hebert PDN, Cywinska A, Ball SL, deWaard JR (2003) Biological identifications through DNA barcodes. Proceedings of the Royal Society of London. Series B: Biological Sciences 270(1512):313–321. doi: 10.1098/rspb.2002.2218.

Liaw A, Wiener M (2002) Classification and Regression by randomForest. R news 2(3):18–22.

McAney CM, Fairley JS (1989) Analysis of the diet of the lesser horseshoe bat Rhinolophus hipposideros in the West of Ireland. Journal of Zoology, 217:491–498.

McDevitt AD, Montgomery WI, Tosh DG, Lusby J, Reid N, White TA, McDevitt CD, O’Halloran J, Searle JB, Yearsley JM (2014) Invading and expanding: Range dynamics and ecological consequences of the greater white-toothed shrew (*Crocidura russula*) invasion in Ireland. PLoS ONE 9(6). doi: 10.1371/journal.pone.0100403.

Meharg MJ, Montgomery WI, Dunwoody T (1990) Trophic Relationships of Common Frog (*Rana temporaria*) and Pygmy Shrew (*Sorex minutus*) in Upland Co Antrim, Northern- Ireland. Journal of Zoology 222:1–17.

Novak T, Csuzdi C, Janžekovič F, Pipan T, Devetak D, Lipovšek S (2014) Survival of the epigean Dendrodrilus rubidus tenuis (Oligochaeta: Lumbricidae) in a subterranean environment. Acta Carsologica 43(2–3):331–338. https://doi.org/10.3986/ac.v43i2.586

Pernetta JC (1976) Diets of the Shrews *Sorex araneus* L. and *Sorex minutus* L. in Wytham Grassland. Journal of Animal Ecology 45(3):899–912.

Piñol J, Mir G, Gomez-Polo P, Agustí N. (2015) Universal and blocking primer mismatches limit the use of high-throughput DNA sequencing for the quantitative metabarcoding of arthropods. Molecular Ecology Resources 15(4):819–830. doi: 10.1111/1755-0998.12355.

Piñol J, Senar MA, Symondson WOC (2018) The choice of universal primers and the characteristics of the species mixture determine when DNA metabarcoding can be quantitative. Molecular Ecology 28(2):407–419. doi: 10.1111/mec.14776.

Pompanon F, Deagle BE, Symondson WOC, Brown DS, Jarman SN, Taberlet P (2012) Who is eating what: Diet assessment using next generation sequencing. Molecular Ecology 21(8):1931–1950. doi: 10.1111/j.1365-294X.2011.05403.x.

Puechmaille S, Mathy G, Petit E (2005) Characterization of 14 polymorphic microsatellite loci for the lesser horseshoe bat, *Rhinolophus hipposideros* (Rhinolophidae, Chiroptera). Molecular Ecology Notes 5:941–944.

Saitoh S, Aoyama H, Fujii S, Sunagawa H, Nagahama H, Akutsu M, Shinzato N, Naneko N, Nakamori T (2016) A quantitative protocol for DNA metabarcoding of springtails (Collembola). The 6th International Barcode of Life Conference. 01:705–723.

Siegenthaler A, Wangensteen OS, Soto AZ, Benvenuto C, Corrigan L, Mariani S (2019) Metabarcoding of shrimp stomach content: Harnessing a natural sampler for fish biodiversity monitoring. Molecular Ecology 19(1):206–220. doi: 10.1111/1755-0998.12956.

Sikes RS (2016) 2016 Guidelines of the American Society of Mammalogists for the use of wild mammals in research and education. Journal of Mammalogy 97(3):663–688. doi: 10.1093/jmammal/gyw078.

Stork NE (2018) How Many Species of Insects and Other Terrestrial Arthropods Are There on Earth?. Annual Review of Entomology 63:31–45.

Tournayre O, Leuchtmann M, Filippi-Cadaccioni O, Trillat M, Piry S, Pontier D, Charbonnel C, Galan M (2020) In silico and empirical evaluation of twelve metabarcoding primer sets for insectivorous diet analyses. Ecology and Evolution, 10:6310–6332. doi: 10.1002/ece3.6362.

Vesterinen EJ, Puisto AIE, Blomberg A, Lilley TM (2018) Table for five, please: Dietary partitioning in boreal bats. Ecology and Evolution 8:10914–10937. doi: 10.1002/ece3.4559.

Ware RL, Booker AL, Allaby FR, Allaby RG (2020) Habitat selection drives dietary specialisation in *Sorex minutus*. bioRxiv. doi: 10.1101/2020.02.03.932913.

Zeale MRK, Butlin RK, Barker GLA, Lees DC, Jones G (2011) Taxon-specific PCR for DNA barcoding arthropod prey in bat faeces. Molecular Ecology Resources 11(2) 236–244. doi: 10.1111/j.1755-0998.2010.02920.x.

Zhang J. (2016) Species Association Analysis “spaa”: R Package.

