## Supplementary Material for "Primer biases in the molecular assessment of diet in multiple insectivorous mammals"

### Appendix 1

#### *Library preparation and sequencing*

To remove unwanted fragments smaller than 100 bp, a left-side bead clean was performed using HighPrep™ PCR Clean-up System (MAGBio Genomics) beads at a 1.1x ratio. To remove fragments larger than 300 bp, a right-side bead clean was performed using a 0.8x ratio of beads to DNA template. Sequencing adapters (Illumina TruSeq dual-index adapters) were ligated onto fragments using the NextFlex PCR-free DNA-seq kit (for Illumina Platforms) following the manufacturer’s protocol. Two sequencing libraries were created; one for the two shrew species combined and one for bats. Library concentration was quantified by qPCR using the NEBnext library quant kit for Illumina (New England Biolabs). Both libraries were pooled at equimolar concentrations and sequenced on a single Illumina MiSeq run using a V2 300 cycle kit with a load concentration of 9 pM and a 1% PhiX spike.

#### *Bioinformatics*

Quality of sequences were examined using FastQC (Andrews 2010). Processing of raw sequence reads to the final dataset was performed using OBITools metabarcoding software (Boyer et al. 2016). High sequence quality required no trimming prior to alignment. Paired end sequences were aligned using *illuminapairedend,* with any reads below a Phred quality score of 40 being discarded. Sequences were demultiplexed using *ngsfilter* according to the unique dual MID tag combination for each sample. Sequences between 128 bp and 138 bp in length from the Gillet primer dataset were retained, while sequences between 152 bp and 162 bp for the Zeale primer dataset were retained. Each library (bat and two shrews) was processed separately up to this point. The demultiplexed fastq files were concatenated so that the bat and two shrew libraries were processed together. Unique sequences were combined using *obiuniq*. Chimeras were detected and removed using the uchime denovo method (Edgar et al. 2011) in VSEARCH (Rognes et al. 2016).

Sequences were clustered into MOTUs using *sumaclust*. Empirically testing different values is recommended (Alberdi et al. 2018) and to explore the number of MOTUs identified and taxonomically assigned, sequences were clustered using a range from 95% to 98%, which are commonly used similarity values for the COI region (Clare et al. 2011; Razgour et al. 2011). Further analyses were processed using the 98% clustering threshold based off empirical results.

Singletons (MOTUs represented by a total of one sequence read in the entire dataset) were removed prior to taxonomic assignment to aid computation time. MOTUs were taxonomically assigned using blastn against the GenBank database. Sequences required at least 80% identity and 90% alignment to match against taxa. The top 25 hits were returned, and the most common taxid was assigned. MOTUs required at least 98% identity for species level assignment (Arrizabalaga-Escudero et al. 2018; Clare et al. 2014). MOTUs between 95% and 98% were restricted to genus level assignment. MOTUs between 93% and 95% were restricted to family level assignment. MOTUs between 90% and 93% were restricted to order level assignment.

Blasting against the GenBank database was used because GenBank has a larger repository than BOLD, and a recent study shows that for insect taxa, GenBank can outperform BOLD for species level identification (Meiklejohn et al. 2019). MOTUs with lower taxonomic resolution were manually cross-referenced using the BOLD repository to increase taxonomic resolution where possible.

*MOTU filtering*

MOTUs of which more than 2% of the reads were found in the blanks were removed. All MOTUs belonging to non-prey taxa (such as vertebrates and parasites), identified using the GenBank database, were removed. Samples with less than 1000 reads were also removed. To avoid the inclusion of false positive taxa, MOTUs were removed from each sample if they were represented by less than 0.01% of the total reads of that individual sample (Alberdi et al. 2018). This is used rather than an absolute threshold to account for variable sequencing depth of samples.

As the Zeale and Gillet primers amplify slightly different regions of COI, sequences were clustered into MOTUs separately. It is possible that the Zeale and Gillet primers may detect two separate MOTUs of unknown species to the same genus level. For example, Primer A detects MOTU-A without the reference sequence to assign to species, but it can be assigned to the genus level. Primer B detects MOTU-B without the reference sequence to assign to species, but it can be assigned to the same genus as MOTU-A. There is no way to definitively determine if both these MOTUs are two different species or the same species split into two MOTUs. The latter case can artificially inflate diversity measures. Consequently, diversity measures between primers, was measured by agglomerating MOTUs to the highest taxonomic resolution available. The results for each library will be presented according to primer; Gillet, Zeale and Both (i.e. both primer datasets combined).

To determine the coverage of samples, rarefaction curves were generated using the R package vegan (Oksanen et al. 2019). In addition, the *depth_cov()* function in the hilldiv R package (Alberdi and Gilbert, 2019) was used to clarify if sufficient read depth was obtained for each sample, using the qvalue = 1 (equivalent to Shannon diversity measure).

All bioinformatic steps and scripts can be found on github (<https://github.com/ShrewlockHolmes>). Raw sequence data will be made publicly available upon publication.

### Tables

Table S1. Sampling locations for *Rhinolophus hipposideros*, *Sorex minutus* and *Crocidura russula* in Ireland and Belle Ile. See Figure S1 for approximate locations.

| **Map no.** | **Species** | **Latitude** | **Longitude** |
| --- | --- | --- | --- |
| 1 | *Rhinolophus hipposideros* | 53.55 | -9.35 |
| 2 | *Rhinolophus hipposideros* | 53.08 | -8.88 |
| 3 | *Rhinolophus hipposideros* | 52.89 | -9.03 |
| 4 | *Rhinolophus hipposideros* | 52.59 | -8.87 |
| 5 | *Rhinolophus hipposideros* | 52.01 | -9.50 |
| 6 | *Rhinolophus hipposideros* | 51.74 | -9.52 |
| 7 | *Sorex minutus* | 52.55 | -9.15 |
| 8 | *Crocidura russula* | 52.62 | -8.70 |
| 9 | *Crocidura russula* | 52.57 | -6.99 |
| 10 | *Sorex minutus + Crocidura russula* | 51.99 | -7.94 |
| 11 | *Sorex minutus + Crocidura russula* | 52.57 | -6.99 |
| 12 | *Sorex minutus + Crocidura russula* | 47.34 | -3.24 |

Table S2. The 20 prey/taxa with the highest MeanDecreaseGini values from the random forest classifier analysis. These taxa have the most influence in differentiating bat and shrew samples when using the Zeale primers

| **MOTU** | **class** | **order** | **family** | **genus** | **species** | **MeanDecreaseGini** |
| --- | --- | --- | --- | --- | --- | --- |
| zMOTU_27 | Insecta | Diptera | Tipulidae | *Tipula* | Genus_*Tipula* | 1.736948676 |
| zMOTU_3 | Insecta | Diptera | Limoniidae | Family_Limoniidae | Family_Limoniidae | 1.208434072 |
| zMOTU_13 | Insecta | Diptera | Tipulidae | Family_Tipulidae | Family_Tipulidae | 1.137379781 |
| zMOTU_4 | Insecta | Lepidoptera | Yponomeutidae | *Prays* | *Prays_fraxinella* | 0.588655306 |
| zMOTU_12 | Insecta | Coleoptera | Tenebrionidae | *Lagria* | *Lagria_hirta* | 0.432887792 |
| zMOTU_59 | Insecta | Diptera | Tipulidae | *Tipula* | *Tipula_furca* | 0.413486862 |
| zMOTU_1 | Insecta | Lepidoptera | Noctuidae | *Lycophotia* | *Lycophotia_porphyrea* | 0.369636471 |
| zMOTU_266 | Insecta | Lepidoptera | Yponomeutidae | *Prays* | Genus_*Prays* | 0.322773668 |
| zMOTU_51 | Diplopoda | Julida | Julidae | *Leptoiulus* | Genus_*Leptoiulus* | 0.313260049 |
| zMOTU_209 | Insecta | Diptera | Limoniidae | Family_Limoniidae | Family_Limoniidae | 0.304900105 |
| zMOTU_17 | Insecta | Lepidoptera | Erebidae | *Spilarctia* | *Spilarctia_luteum* | 0.298069174 |
| zMOTU_2 | Insecta | Diptera | Order_Diptera | Order_Diptera | Order_Diptera | 0.240581244 |
| zMOTU_18 | Insecta | Diptera | Tipulidae | *Tipula* | *Tipula_oleracea* | 0.224616968 |
| zMOTU_65 | Insecta | Trichoptera | Limnephilidae | *Limnephilus* | *Limnephilus_stigma* | 0.222774098 |
| gMOTU_88 | Insecta | Lepidoptera | Hepialidae | *Pharmacis* | *Pharmacis_fusconebulosa* | 0.208164351 |
| zMOTU_494 | Insecta | Coleoptera | Tenebrionidae | *Lagria* | Genus_*Lagria* | 0.204610293 |
| zMOTU_300 | Insecta | Coleoptera | Staphylinidae | *Quedius* | Genus_*Quedius* | 0.188613613 |
| zMOTU_99 | Insecta | Diptera | Bibionidae | Family_Bibionidae | Family_Bibionidae | 0.188020667 |
| zMOTU_133 | Insecta | Coleoptera | Staphylinidae | Family_Staphylinidae | Family_Staphylinidae | 0.185003148 |
| zMOTU_156 | Insecta | Diptera | Cecidomyiidae | Family_Cecidomyiidae | Family_Cecidomyiidae | 0.17802937 |

Table S3. The 20 prey/taxa with the highest MeanDecreaseGini values from the random forest classifier analysis. These taxa have the most influence in differentiating bat and shrew samples when using the Gillet Primers

| **MOTU** | **class** | **order** | **family** | **genus** | **species** | **MeanDecreaseGini** |
| --- | --- | --- | --- | --- | --- | --- |
| gMOTU_34 | Insecta | Diptera | Tipulidae | *Tipula* | Genus_*Tipula* | 1.548673533 |
| gMOTU_51 | Insecta | Diptera | Limoniidae | *Austrolimnophila* | *Austrolimnophila_ochracea* | 0.957704144 |
| gMOTU_276 | Insecta | Diptera | Cecidomyiidae | Family_Cecidomyiidae | Family_Cecidomyiidae | 0.902665919 |
| gMOTU_36 | Insecta | Diptera | Tipulidae | Family_Tipulidae | Family_Tipulidae | 0.876208733 |
| gMOTU_60 | Insecta | Diptera | Order_Diptera | Order_Diptera | Order_Diptera | 0.776165915 |
| gMOTU_66 | Insecta | Lepidoptera | Yponomeutidae | *Prays* | *Prays_oleae* | 0.707225936 |
| gMOTU_71 | Insecta | Diptera | Agromyzidae | Family_Agromyzidae | Family_Agromyzidae | 0.662690674 |
| zMOTU_65 | Insecta | Trichoptera | Limnephilidae | *Limnephilus* | *Limnephilus_stigma* | 0.571967754 |
| zMOTU_59 | Insecta | Diptera | Tipulidae | *Tipula* | *Tipula_furca* | 0.538942691 |
| gMOTU_300 | Insecta | Trichoptera | Limnephilidae | *Limnephilus* | Genus_*Limnephilus* | 0.385366204 |
| gMOTU_48 | Arachnida | Class_Arachnida | Class_Arachnida | Class_Arachnida | Class_Arachnida | 0.350730148 |
| gMOTU_40 | Insecta | Diptera | Tipulidae | *Tipula* | *Tipula_parshleyi* | 0.335058742 |
| gMOTU_135 | Insecta | Diptera | Tephritidae | Family_Tephritidae | Family_Tephritidae | 0.314000321 |
| gMOTU_130 | Insecta | Diptera | Limoniidae | *Neolimnophila* | Genus_*Neolimnophila* | 0.313094263 |
| gMOTU_14 | Insecta | Diptera | Limoniidae | *Molophilus* | *Molophilus_griseus* | 0.293392929 |
| gMOTU_84 | Insecta | Class_Insecta | Class_Insecta | Class_Insecta | Class_Insecta | 0.272686585 |
| zMOTU_81 | Insecta | Lepidoptera | Tortricidae | *Pseudargyrotoza* | *Pseudargyrotoza_conwagana* | 0.258685263 |
| zMOTU_8 | Insecta | Coleoptera | Curculionidae | *Caenopsis* | *Caenopsis_waltoni* | 0.249718179 |
| gMOTU_20 | Insecta | Diptera | Tipulidae | *Tipula* | Genus_*Tipula* | 0.245305575 |
| gMOTU_155 | Insecta | Diptera | Scathophagidae | *Scathophaga* | Genus_*Scathophaga* | 0.243618028 |

Table S4. The 20 prey/taxa with the highest MeanDecreaseGini values from the random forest classifier analysis. These taxa have the most influence in differentiating bat and shrew samples when using Both Primers

| **MOTU** | **class** | **order** | **family** | **genus** | **species** | **MeanDecreaseGini** |
| --- | --- | --- | --- | --- | --- | --- |
| zMOTU_27 | Insecta | Diptera | Tipulidae | *Tipula* | Genus_*Tipula* | 1.888741201 |
| zMOTU_13 | Insecta | Diptera | Tipulidae | Family_Tipulidae | Family_Tipulidae | 1.236129877 |
| zMOTU_3 | Insecta | Diptera | Limoniidae | Family_Limoniidae | Family_Limoniidae | 1.146088842 |
| zMOTU_156 | Insecta | Diptera | Cecidomyiidae | Family_Cecidomyiidae | Family_Cecidomyiidae | 0.939196938 |
| gMOTU_51 | Insecta | Diptera | Limoniidae | *Austrolimnophila* | *Austrolimnophila_ochracea* | 0.692626856 |
| zMOTU_59 | Insecta | Diptera | Tipulidae | *Tipula* | *Tipula_furca* | 0.664268419 |
| zMOTU_65 | Insecta | Trichoptera | Limnephilidae | *Limnephilus* | *Limnephilus_stigma* | 0.575748805 |
| zMOTU_4 | Insecta | Lepidoptera | Yponomeutidae | *Prays* | *Prays_fraxinella* | 0.563766435 |
| gMOTU_66 | Insecta | Lepidoptera | Yponomeutidae | *Prays* | *Prays_oleae* | 0.419467486 |
| zMOTU_266 | Insecta | Lepidoptera | Yponomeutidae | *Prays* | Genus_*Prays* | 0.418180229 |
| gMOTU_71 | Insecta | Diptera | Agromyzidae | Family_Agromyzidae | Family_Agromyzidae | 0.360473597 |
| zMOTU_2 | Insecta | Diptera | Order_Diptera | Order_Diptera | Order_Diptera | 0.313490984 |
| gMOTU_48 | Arachnida | Class_Arachnida | Class_Arachnida | Class_Arachnida | Class_Arachnida | 0.291101537 |
| gMOTU_300 | Insecta | Trichoptera | Limnephilidae | *Limnephilus* | Genus_*Limnephilus* | 0.277312333 |
| gMOTU_14 | Insecta | Diptera | Limoniidae | *Molophilus* | *Molophilus_griseus* | 0.256437941 |
| gMOTU_88 | Insecta | Lepidoptera | Hepialidae | *Pharmacis* | *Pharmacis_fusconebulosa* | 0.254544787 |
| zMOTU_18 | Insecta | Diptera | Tipulidae | *Tipula* | *Tipula_oleracea* | 0.244565058 |
| zMOTU_81 | Insecta | Lepidoptera | Tortricidae | *Pseudargyrotoza* | *Pseudargyrotoza_conwagana* | 0.231654187 |
| zMOTU_8 | Insecta | Coleoptera | Curculionidae | *Caenopsis* | *Caenopsis_waltoni* | 0.215343719 |
| zMOTU_209 | Insecta | Diptera | Limoniidae | Family_Limoniidae | Family_Limoniidae | 0.209839028 |

Table S5. The 20 prey/taxa with the highest MeanDecreaseGini values from the random forest classifier analysis. These taxa have the most influence in differentiating *C. russula* and *S. minutus* samples when using the Zeale Primers

| **MOTU** | **class** | **order** | **family** | **genus** | **species** | **MeanDecreaseGini** |
| --- | --- | --- | --- | --- | --- | --- |
| zMOTU_33 | Insecta | Lepidoptera | Noctuidae | *Hoplodrina* | *Hoplodrina_blanda* | 0.210364547 |
| zMOTU_253 | Insecta | Class_Insecta | Class_Insecta | Class_Insecta | Class_Insecta | 0.204989491 |
| zMOTU_12 | Insecta | Coleoptera | Tenebrionidae | *Lagria* | *Lagria_hirta* | 0.184111763 |
| zMOTU_968 | Insecta | Lepidoptera | Erebidae | *Spilarctia* | Genus_*Spilarctia* | 0.15758544 |
| zMOTU_11 | Insecta | Coleoptera | Staphylinidae | *Quedius* | *Quedius_fuliginosus* | 0.11594482 |
| zMOTU_51 | Diplopoda | Julida | Julidae | *Leptoiulus* | Genus_*Leptoiulus* | 0.114844354 |
| zMOTU_524 | Insecta | Lepidoptera | Xyloryctidae | *Cryptophasa* | Genus_*Cryptophasa* | 0.110975386 |
| zMOTU_109 | Insecta | Coleoptera | Carabidae | Family_Carabidae | Family_Carabidae | 0.106080639 |
| zMOTU_1257 | Insecta | Lepidoptera | Glyphipterigidae | Family_Glyphipterigidae | Family_Glyphipterigidae | 0.104970721 |
| zMOTU_17 | Insecta | Lepidoptera | Erebidae | *Spilarctia* | *Spilarctia_luteum* | 0.103195472 |
| zMOTU_2 | Insecta | Diptera | Order_Diptera | Order_Diptera | Order_Diptera | 0.102600392 |
| zMOTU_603 | Insecta | Coleoptera | Carabidae | *Pterostichus* | Genus_*Pterostichus* | 0.102241825 |
| zMOTU_329 | Insecta | Hymenoptera | Order_Hymenoptera | Order_Hymenoptera | Order_Hymenoptera | 0.099288462 |
| zMOTU_46 | Insecta | Coleoptera | Carabidae | *Pterostichus* | *Pterostichus_madidus* | 0.098279377 |
| zMOTU_570 | Insecta | Lepidoptera | Noctuidae | *Hoplodrina* | Genus_*Hoplodrina* | 0.096591615 |
| zMOTU_1120 | Diplopoda | Julida | Julidae | Family_Julidae | Family_Julidae | 0.095153255 |
| zMOTU_402 | Insecta | Lepidoptera | Pyralidae | *Endotricha* | *Endotricha_flammealis* | 0.093672966 |
| zMOTU_377 | Insecta | Lepidoptera | Noctuidae | *Caradrina* | *Caradrina_fuscicornis* | 0.089630734 |
| zMOTU_300 | Insecta | Coleoptera | Staphylinidae | *Quedius* | Genus_*Quedius* | 0.089602765 |
| zMOTU_1860 | Insecta | Diptera | Lonchopteridae | *Lonchoptera* | Genus_*Lonchoptera* | 0.080817573 |

Table S6. The 20 prey/taxa with the highest MeanDecreaseGini values from the random forest classifier analysis. These taxa have the most influence in differentiating *C. russula* and *S. minutus* samples when using the Gillet Primers

| **MOTU** | **class** | **order** | **family** | **genus** | **species** | **MeanDecreaseGini** |
| --- | --- | --- | --- | --- | --- | --- |
| gMOTU_98 | Clitellata | Haplotaxida | Lumbricidae | *Satchellius* | *Satchellius_mammalis* | 0.454919545 |
| gMOTU_234 | Clitellata | Haplotaxida | Lumbricidae | Family_Lumbricidae | Family_Lumbricidae | 0.298436765 |
| gMOTU_75 | Arachnida | Opiliones | Leiobunidae | *Leiobunum* | *Leiobunum_blackwalli* | 0.282072081 |
| gMOTU_28 | Diplopoda | Polydesmida | Polydesmidae | *Polydesmus* | Genus_*Polydesmus* | 0.276982359 |
| gMOTU_983 | Insecta | Diptera | Limoniidae | Family_Limoniidae | Family_Limoniidae | 0.245044277 |
| gMOTU_44 | Insecta | Hymenoptera | Braconidae | *Microplitis* | Genus_*Microplitis* | 0.222259759 |
| gMOTU_84 | Insecta | Class_Insecta | Class_Insecta | Class_Insecta | Class_Insecta | 0.186067386 |
| gMOTU_48 | Arachnida | Class_Arachnida | Class_Arachnida | Class_Arachnida | Class_Arachnida | 0.184524083 |
| gMOTU_38 | Clitellata | Haplotaxida | Lumbricidae | *Lumbricus* | *Lumbricus_castaneus* | 0.184369204 |
| gMOTU_5 | Malacostraca | Isopoda | Armadillidiidae | *Armadillidium* | *Armadillidium_vulgare* | 0.170796895 |
| gMOTU_166 | Gastropoda | Stylommatophora | Agriolimacidae | *Deroceras* | *Deroceras_reticulatum* | 0.167210119 |
| gMOTU_153 | Diplopoda | Julida | Julidae | *Cylindroiulus* | *Cylindroiulus_latestriatus* | 0.158297025 |
| gMOTU_14 | Insecta | Diptera | Limoniidae | *Molophilus* | *Molophilus_griseus* | 0.15058316 |
| zMOTU_11 | Insecta | Coleoptera | Staphylinidae | *Quedius* | *Quedius_fuliginosus* | 0.145617879 |
| gMOTU_82 | Arachnida | Mesostigmata | Order_Mesostigmata | Order_Mesostigmata | Order_Mesostigmata | 0.144629296 |
| gMOTU_986 | Diplopoda | Julida | Julidae | *Ophyiulus* | *Ophyiulus_pilosus* | 0.141491183 |
| gMOTU_13 | Gastropoda | Stylommatophora | Agriolimacidae | *Deroceras* | *Deroceras_laeve* | 0.141381478 |
| gMOTU_788 | Arachnida | Opiliones | Leiobunidae | *Leiobunum* | Genus_*Leiobunum* | 0.140165744 |
| gMOTU_1373 | Collembola | Entomobryomorpha | Tomoceridae | *Tomocerus* | Genus_*Tomocerus* | 0.135909069 |
| gMOTU_60 | Insecta | Diptera | Order_Diptera | Order_Diptera | Order_Diptera | 0.135434044 |

Table S7. The 20 prey/taxa with the highest MeanDecreaseGini values from the random forest classifier analysis. These taxa have the most influence in differentiating *C. russula* and *S. minutus* samples when using Both Primers

| **MOTU** | **class** | **order** | **family** | **genus** | **species** | **MeanDecreaseGini** |
| --- | --- | --- | --- | --- | --- | --- |
| gMOTU_98 | Clitellata | Haplotaxida | Lumbricidae | *Satchellius* | *Satchellius_mammalis* | 0.462844623 |
| gMOTU_234 | Clitellata | Haplotaxida | Lumbricidae | Family_Lumbricidae | Family_Lumbricidae | 0.288619524 |
| gMOTU_75 | Arachnida | Opiliones | Leiobunidae | *Leiobunum* | *Leiobunum_blackwalli* | 0.28289758 |
| gMOTU_28 | Diplopoda | Polydesmida | Polydesmidae | *Polydesmus* | Genus_*Polydesmus* | 0.242112485 |
| gMOTU_84 | Insecta | Class_Insecta | Class_Insecta | Class_Insecta | Class_Insecta | 0.228826433 |
| gMOTU_983 | Insecta | Diptera | Limoniidae | Family_Limoniidae | Family_Limoniidae | 0.215004402 |
| gMOTU_44 | Insecta | Hymenoptera | Braconidae | *Microplitis* | Genus_*Microplitis* | 0.211994962 |
| gMOTU_38 | Clitellata | Haplotaxida | Lumbricidae | *Lumbricus* | *Lumbricus_castaneus* | 0.189272497 |
| gMOTU_153 | Diplopoda | Julida | Julidae | *Cylindroiulus* | *Cylindroiulus_latestriatus* | 0.152445099 |
| gMOTU_986 | Diplopoda | Julida | Julidae | *Ophyiulus* | *Ophyiulus_pilosus* | 0.147364149 |
| gMOTU_5 | Malacostraca | Isopoda | Armadillidiidae | *Armadillidium* | *Armadillidium_vulgare* | 0.146939052 |
| gMOTU_82 | Arachnida | Mesostigmata | Order_Mesostigmata | Order_Mesostigmata | Order_Mesostigmata | 0.138636108 |
| zMOTU_2 | Insecta | Diptera | Order_Diptera | Order_Diptera | Order_Diptera | 0.138567128 |
| gMOTU_13 | Gastropoda | Stylommatophora | Agriolimacidae | *Deroceras* | *Deroceras_laeve* | 0.134356417 |
| gMOTU_14 | Insecta | Diptera | Limoniidae | *Molophilus* | *Molophilus_griseus* | 0.131849295 |
| gMOTU_48 | Arachnida | Class_Arachnida | Class_Arachnida | Class_Arachnida | Class_Arachnida | 0.131132341 |
| gMOTU_788 | Arachnida | Opiliones | Leiobunidae | *Leiobunum* | Genus_*Leiobunum* | 0.128294221 |
| zMOTU_11 | Insecta | Coleoptera | Staphylinidae | *Quedius* | Quedius_fuliginosus | 0.124886427 |
| gMOTU_1373 | Collembola | Entomobryomorpha | Tomoceridae | *Tomocerus* | Genus_*Tomocerus* | 0.124396441 |
| gMOTU_71 | Insecta | Diptera | Agromyzidae | Family_Agromyzidae | Family_Agromyzidae | 0.120293987 |

Table S8. The 20 prey/taxa with the highest MeanDecreaseGini values from the random forest classifier analysis. These taxa have the most influence in differentiating *R. hipposideros* samples amplified with the Zeale or Gillet Primers

| **MOTU** | **class** | **order** | **family** | **genus** | **species** | **MeanDecreaseGini** |
| --- | --- | --- | --- | --- | --- | --- |
| zMOTU_3 | Insecta | Diptera | Limoniidae | Family_Limoniidae | Family_Limoniidae | 1.331839457 |
| gMOTU_51 | Insecta | Diptera | Limoniidae | *Austrolimnophila* | *Austrolimnophila_ochracea* | 1.27655919 |
| gMOTU_71 | Insecta | Diptera | Agromyzidae | Family_Agromyzidae | Family_Agromyzidae | 1.086106311 |
| gMOTU_20 | Insecta | Diptera | Tipulidae | *Tipula* | Genus_*Tipula* | 0.915576631 |
| gMOTU_66 | Insecta | Lepidoptera | Yponomeutidae | *Prays* | *Prays_oleae* | 0.740683569 |
| zMOTU_201 | Insecta | Class_Insecta | Class_Insecta | Class_Insecta | Class_Insecta | 0.691327401 |
| zMOTU_4 | Insecta | Lepidoptera | Yponomeutidae | *Prays* | *Prays_fraxinella* | 0.568687875 |
| gMOTU_300 | Insecta | Trichoptera | Limnephilidae | *Limnephilus* | Genus_*Limnephilus* | 0.492781379 |
| zMOTU_266 | Insecta | Lepidoptera | Yponomeutidae | *Prays* | Genus_*Prays* | 0.418688161 |
| gMOTU_55 | Insecta | Diptera | Bibionidae | Family_Bibionidae | Family_Bibionidae | 0.376624294 |
| gMOTU_40 | Insecta | Diptera | Tipulidae | *Tipula* | *Tipula_parshleyi* | 0.356506672 |
| zMOTU_13 | Insecta | Diptera | Tipulidae | Family_Tipulidae | Family_Tipulidae | 0.355780051 |
| gMOTU_130 | Insecta | Diptera | Limoniidae | *Neolimnophila* | Genus_*Neolimnophila* | 0.353903434 |
| gMOTU_135 | Insecta | Diptera | Tephritidae | Family_Tephritidae | Family_Tephritidae | 0.346765019 |
| gMOTU_117 | Insecta | Neuroptera | Hemerobiidae | *Hemerobius* | *Hemerobius_humulinus* | 0.333296852 |
| zMOTU_27 | Insecta | Diptera | Tipulidae | *Tipula* | Genus_*Tipula* | 0.326350712 |
| zMOTU_209 | Insecta | Diptera | Limoniidae | Family_Limoniidae | Family_Limoniidae | 0.319077622 |
| gMOTU_155 | Insecta | Diptera | Scathophagidae | *Scathophaga* | Genus_*Scathophaga* | 0.265024661 |
| zMOTU_42 | Insecta | Diptera | Order_Diptera | Order_Diptera | Order_Diptera | 0.257537177 |
| zMOTU_156 | Insecta | Diptera | Cecidomyiidae | Family_Cecidomyiidae | Family_Cecidomyiidae | 0.254402663 |

Table S9. The 20 prey/taxa with the highest MeanDecreaseGini values from the random forest classifier analysis. These taxa have the most influence in differentiating *R. hipposideros* samples amplified with the Zeale or Both Primers

| **MOTU** | **class** | **order** | **family** | **genus** | **species** | **MeanDecreaseGini** |
| --- | --- | --- | --- | --- | --- | --- |
| gMOTU_51 | Insecta | Diptera | Limoniidae | *Austrolimnophila* | *Austrolimnophila_ochracea* | 1.319704373 |
| gMOTU_20 | Insecta | Diptera | Tipulidae | *Tipula* | Genus_Tipula | 0.953832406 |
| gMOTU_66 | Insecta | Lepidoptera | Yponomeutidae | *Prays* | *Prays_oleae* | 0.804693942 |
| gMOTU_71 | Insecta | Diptera | Agromyzidae | Family_Agromyzidae | Family_Agromyzidae | 0.77519654 |
| zMOTU_201 | Insecta | Class_Insecta | Class_Insecta | Class_Insecta | Class_Insecta | 0.539585075 |
| gMOTU_40 | Insecta | Diptera | Tipulidae | *Tipula* | *Tipula_parshleyi* | 0.441981955 |
| gMOTU_130 | Insecta | Diptera | Limoniidae | *Neolimnophila* | Genus_*Neolimnophila* | 0.399654702 |
| gMOTU_117 | Insecta | Neuroptera | Hemerobiidae | *Hemerobius* | *Hemerobius_humulinus* | 0.381504238 |
| gMOTU_135 | Insecta | Diptera | Tephritidae | Family_Tephritidae | Family_Tephritidae | 0.377640997 |
| gMOTU_55 | Insecta | Diptera | Bibionidae | Family_Bibionidae | Family_Bibionidae | 0.377026711 |
| gMOTU_300 | Insecta | Trichoptera | Limnephilidae | *Limnephilus* | Genus_*Limnephilus* | 0.34881122 |
| zMOTU_156 | Insecta | Diptera | Cecidomyiidae | Family_Cecidomyiidae | Family_Cecidomyiidae | 0.332484935 |
| gMOTU_155 | Insecta | Diptera | Scathophagidae | *Scathophaga* | Genus_*Scathophaga* | 0.303359724 |
| zMOTU_42 | Insecta | Diptera | Order_Diptera | Order_Diptera | Order_Diptera | 0.281455043 |
| zMOTU_3 | Insecta | Diptera | Limoniidae | Family_Limoniidae | Family_Limoniidae | 0.278798297 |
| zMOTU_27 | Insecta | Diptera | Tipulidae | *Tipula* | Genus_*Tipula* | 0.278477145 |
| zMOTU_13 | Insecta | Diptera | Tipulidae | Family_Tipulidae | Family_Tipulidae | 0.271373602 |
| zMOTU_18 | Insecta | Diptera | Tipulidae | *Tipula* | *Tipula_oleracea* | 0.228999393 |
| gMOTU_218 | Insecta | Diptera | Cecidomyiidae | *Macrodiplosis* | Genus_*Macrodiplosis* | 0.228454199 |
| gMOTU_241 | Insecta | Lepidoptera | Tortricidae | *Celypha* | *Celypha_lacunana* | 0.211379436 |

Table S10. The 20 prey/taxa with the highest MeanDecreaseGini values from the random forest classifier analysis. These taxa have the most influence in differentiating *R. hipposideros* samples amplified with the Gillet or Both Primers

| **MOTU** | **class** | **order** | **family** | **genus** | **species** | **MeanDecreaseGini** |
| --- | --- | --- | --- | --- | --- | --- |
| zMOTU_3 | Insecta | Diptera | Limoniidae | Family_Limoniidae | Family_Limoniidae | 1.438880029 |
| gMOTU_51 | Insecta | Diptera | Limoniidae | *Austrolimnophila* | *Austrolimnophila_ochracea* | 0.748043213 |
| zMOTU_4 | Insecta | Lepidoptera | Yponomeutidae | *Prays* | *Prays_fraxinella* | 0.691359217 |
| zMOTU_201 | Insecta | Class_Insecta | Class_Insecta | Class_Insecta | Class_Insecta | 0.538220101 |
| gMOTU_71 | Insecta | Diptera | Agromyzidae | Family_Agromyzidae | Family_Agromyzidae | 0.515947776 |
| gMOTU_20 | Insecta | Diptera | Tipulidae | *Tipula* | Genus_*Tipula* | 0.497570944 |
| gMOTU_66 | Insecta | Lepidoptera | Yponomeutidae | *Prays* | *Prays_oleae* | 0.482221401 |
| zMOTU_266 | Insecta | Lepidoptera | Yponomeutidae | *Prays* | Genus_*Prays* | 0.480059006 |
| zMOTU_13 | Insecta | Diptera | Tipulidae | Family_Tipulidae | Family_Tipulidae | 0.456014824 |
| zMOTU_209 | Insecta | Diptera | Limoniidae | Family_Limoniidae | Family_Limoniidae | 0.407264496 |
| zMOTU_27 | Insecta | Diptera | Tipulidae | *Tipula* | Genus_*Tipula* | 0.365104021 |
| gMOTU_135 | Insecta | Diptera | Tephritidae | Family_Tephritidae | Family_Tephritidae | 0.330022286 |
| zMOTU_42 | Insecta | Diptera | Order_Diptera | Order_Diptera | Order_Diptera | 0.280162318 |
| gMOTU_117 | Insecta | Neuroptera | Hemerobiidae | *Hemerobius* | *Hemerobius_humulinus* | 0.275703448 |
| gMOTU_185 | Insecta | Diptera | Tipulidae | *Tipula* | *Tipula_banffiana* | 0.249166959 |
| zMOTU_65 | Insecta | Trichoptera | Limnephilidae | *Limnephilus* | *Limnephilus_stigma* | 0.246238084 |
| zMOTU_227 | Insecta | Diptera | Tipulidae | Family_ Tipulidae | Family_ Tipulidae | 0.244443102 |
| zMOTU_55 | Insecta | Diptera | Culicidae | *Culex* | *Culex_pipiens* | 0.228739319 |
| gMOTU_300 | Insecta | Trichoptera | Limnephilidae | *Limnephilus* | Genus_*Limnephilus* | 0.225236701 |
| zMOTU_156 | Insecta | Diptera | Cecidomyiidae | Family_Cecidomyiidae | Family_Cecidomyiidae | 0.223557504 |

Table S11. The 20 prey/taxa with the highest MeanDecreaseGini values from the random forest classifier analysis. These taxa have the most influence in differentiating shrew samples amplified with the Zeale or Gillet Primers

| **MOTU** | **class** | **order** | **family** | **genus** | **species** | **MeanDecreaseGini** |
| --- | --- | --- | --- | --- | --- | --- |
| gMOTU_14 | Insecta | Diptera | Limoniidae | *Molophilus* | *Molophilus_griseus* | 0.67561264 |
| zMOTU_12 | Insecta | Coleoptera | Tenebrionidae | *Lagria* | *Lagria_hirta* | 0.607411095 |
| gMOTU_48 | Arachnida | Class_Arachnida | Class_Arachnida | Class_Arachnida | Class_Arachnida | 0.529790916 |
| zMOTU_51 | Diplopoda | Julida | Julidae | *Leptoiulus* | Genus_*Leptoiulus* | 0.521617062 |
| zMOTU_17 | Insecta | Lepidoptera | Erebidae | *Spilarctia* | *Spilarctia_luteum* | 0.474094375 |
| zMOTU_2 | Insecta | Diptera | Order_Diptera | Order_Diptera | Order_Diptera | 0.359390439 |
| zMOTU_494 | Insecta | Coleoptera | Tenebrionidae | *Lagria* | Genus_*Lagria* | 0.32200955 |
| gMOTU_84 | Insecta | Class_Insecta | Class_Insecta | Class_Insecta | Class_Insecta | 0.302388913 |
| zMOTU_1 | Insecta | Lepidoptera | Noctuidae | *Lycophotia* | *Lycophotia_porphyrea* | 0.275329999 |
| gMOTU_20 | Insecta | Diptera | Tipulidae | *Tipula* | Genus_Tipula | 0.268037508 |
| gMOTU_49 | Insecta | Hemiptera | Delphacidae | Family_Delphacidae | Family_Delphacidae | 0.248368441 |
| gMOTU_93 | Collembola | Entomobryomorpha | Tomoceridae | *Tomocerus* | *Tomocerus_vulgaris* | 0.224474327 |
| gMOTU_5 | Malacostraca | Isopoda | Armadillidiidae | *Armadillidium* | *Armadillidium_vulgare* | 0.214862055 |
| gMOTU_89 | Insecta | Hemiptera | Rhyparochromidae | *Peritrechus* | *Peritrechus_geniculatus* | 0.202953231 |
| zMOTU_11 | Insecta | Coleoptera | Staphylinidae | *Quedius* | *Quedius_fuliginosus* | 0.190641021 |
| zMOTU_8 | Insecta | Coleoptera | Curculionidae | *Caenopsis* | *Caenopsis_waltoni* | 0.166968026 |
| gMOTU_30 | Arachnida | Opiliones | Phalangiidae | *Paroligolophus* | *Paroligolophus_agrestis* | 0.164529235 |
| zMOTU_133 | Insecta | Coleoptera | Staphylinidae | Family_Staphylinidae | Family_Staphylinidae | 0.161878224 |
| gMOTU_55 | Insecta | Diptera | Bibionidae | Family_Bibionidae | Family_Bibionidae | 0.156123145 |
| gMOTU_112 | Insecta | Coleoptera | Staphylinidae | *Quedius* | Genus_*Quedius* | 0.153828673 |

Table S12. The 20 prey/taxa with the highest MeanDecreaseGini values from the random forest classifier analysis. These taxa have the most influence in differentiating shrew samples amplified with the Zeale or Both Primers

| **MOTU** | **class** | **order** | **family** | **genus** | **species** | **MeanDecreaseGini** |
| --- | --- | --- | --- | --- | --- | --- |
| gMOTU_14 | Insecta | Diptera | Limoniidae | *Molophilus* | *Molophilus_griseus* | 0.751677028 |
| gMOTU_48 | Arachnida | Class_Arachnida | Class_Arachnida | Class_Arachnida | Class_Arachnida | 0.616098446 |
| zMOTU_17 | Insecta | Lepidoptera | Erebidae | *Spilarctia* | *Spilarctia_luteum* | 0.344947327 |
| gMOTU_84 | Insecta | Class_Insecta | Class_Insecta | Class_Insecta | Class_Insecta | 0.331007214 |
| zMOTU_2 | Insecta | Diptera | Order_Diptera | Order_Diptera | Order_Diptera | 0.322858133 |
| gMOTU_20 | Insecta | Diptera | Tipulidae | *Tipula* | Genus_*Tipula* | 0.300461807 |
| zMOTU_51 | Diplopoda | Julida | Julidae | *Leptoiulus* | Genus_*Leptoiulus* | 0.299030137 |
| zMOTU_12 | Insecta | Coleoptera | Tenebrionidae | *Lagria* | *Lagria_hirta* | 0.296049576 |
| zMOTU_494 | Insecta | Coleoptera | Tenebrionidae | *Lagria* | *Genus_Lagria* | 0.274148892 |
| gMOTU_5 | Malacostraca | Isopoda | Armadillidiidae | *Armadillidium* | *Armadillidium_vulgare* | 0.252383935 |
| gMOTU_93 | Collembola | Entomobryomorpha | Tomoceridae | *Tomocerus* | *Tomocerus_vulgaris* | 0.252096248 |
| gMOTU_49 | Insecta | Hemiptera | Delphacidae | Family_Delphacidae | Family_Delphacidae | 0.250103075 |
| zMOTU_1 | Insecta | Lepidoptera | Noctuidae | *Lycophotia* | *Lycophotia_porphyrea* | 0.225695628 |
| gMOTU_89 | Insecta | Hemiptera | Rhyparochromidae | *Peritrechus* | *Peritrechus_geniculatus* | 0.213093612 |
| zMOTU_11 | Insecta | Coleoptera | Staphylinidae | *Quedius* | *Quedius_fuliginosus* | 0.191319904 |
| gMOTU_30 | Arachnida | Opiliones | Phalangiidae | *Paroligolophus* | *Paroligolophus_agrestis* | 0.186484073 |
| zMOTU_8 | Insecta | Coleoptera | Curculionidae | *Caenopsis* | *Caenopsis_waltoni* | 0.183868177 |
| zMOTU_268 | Malacostraca | Class_Malacostraca | Class_Malacostraca | Class_Malacostraca | Class_Malacostraca | 0.163846362 |
| gMOTU_112 | Insecta | Coleoptera | Staphylinidae | *Quedius* | Genus_*Quedius* | 0.160627778 |
| gMOTU_46 | Malacostraca | Isopoda | Trichoniscidae | *Trichoniscus* | *Trichoniscus_pusillus* | 0.15664154 |

Table S13. The 20 prey/taxa with the highest MeanDecreaseGini values from the random forest classifier analysis. These taxa have the most influence in differentiating shrew samples amplified with the Gillet or Both Primers

| **MOTU** | **class** | **order** | **family** | **genus** | **species** | **MeanDecreaseGini** |
| --- | --- | --- | --- | --- | --- | --- |
| zMOTU_2 | Insecta | Diptera | Order_Diptera | Order_Diptera | Order_Diptera | 0.446095705 |
| gMOTU_84 | Insecta | Class_Insecta | Class_Insecta | Class_Insecta | Class_Insecta | 0.385783028 |
| zMOTU_12 | Insecta | Coleoptera | Tenebrionidae | *Lagria* | *Lagria_hirta* | 0.383086505 |
| gMOTU_48 | Arachnida | Class_Arachnida | Class_Arachnida | Class_Arachnida | Class_Arachnida | 0.32576506 |
| zMOTU_17 | Insecta | Lepidoptera | Erebidae | *Spilarctia* | *Spilarctia_luteum* | 0.320120538 |
| zMOTU_51 | Diplopoda | Julida | Julidae | *Leptoiulus* | Genus_*Leptoiulus* | 0.315853124 |
| zMOTU_8 | Insecta | Coleoptera | Curculionidae | *Caenopsis* | *Caenopsis_waltoni* | 0.305821471 |
| gMOTU_14 | Insecta | Diptera | Limoniidae | *Molophilus* | *Molophilus_griseus* | 0.2881507 |
| gMOTU_20 | Insecta | Diptera | Tipulidae | *Tipula* | Genus_*Tipula* | 0.260636298 |
| zMOTU_268 | Malacostraca | Class_Malacostraca | Class_Malacostraca | Class_Malacostraca | Class_Malacostraca | 0.251046932 |
| gMOTU_89 | Insecta | Hemiptera | Rhyparochromidae | *Peritrechus* | *Peritrechus_geniculatus* | 0.243388062 |
| gMOTU_49 | Insecta | Hemiptera | Delphacidae | Family_Delphacidae | Family_Delphacidae | 0.232456578 |
| zMOTU_494 | Insecta | Coleoptera | Tenebrionidae | *Lagria* | Genus_*Lagria* | 0.218399925 |
| zMOTU_1 | Insecta | Lepidoptera | Noctuidae | *Lycophotia* | *Lycophotia_porphyrea* | 0.205663988 |
| gMOTU_93 | Collembola | Entomobryomorpha | Tomoceridae | *Tomocerus* | *Tomocerus_vulgaris* | 0.198774184 |
| gMOTU_5 | Malacostraca | Isopoda | Armadillidiidae | *Armadillidium* | *Armadillidium_vulgare* | 0.189881308 |
| gMOTU_46 | Malacostraca | Isopoda | Trichoniscidae | *Trichoniscus* | *Trichoniscus_pusillus* | 0.189517092 |
| gMOTU_13 | Gastropoda | Stylommatophora | Agriolimacidae | *Deroceras* | *Deroceras_laeve* | 0.166971614 |
| gMOTU_31 | Diplopoda | Julida | Order_Julida | Order_Julida | Order_Julida | 0.165763891 |
| gMOTU_161 | Chilopoda | Class_Chilopoda | Class_Chilopoda | Class_Chilopoda | Class_Chilopoda | 0.163469608 |

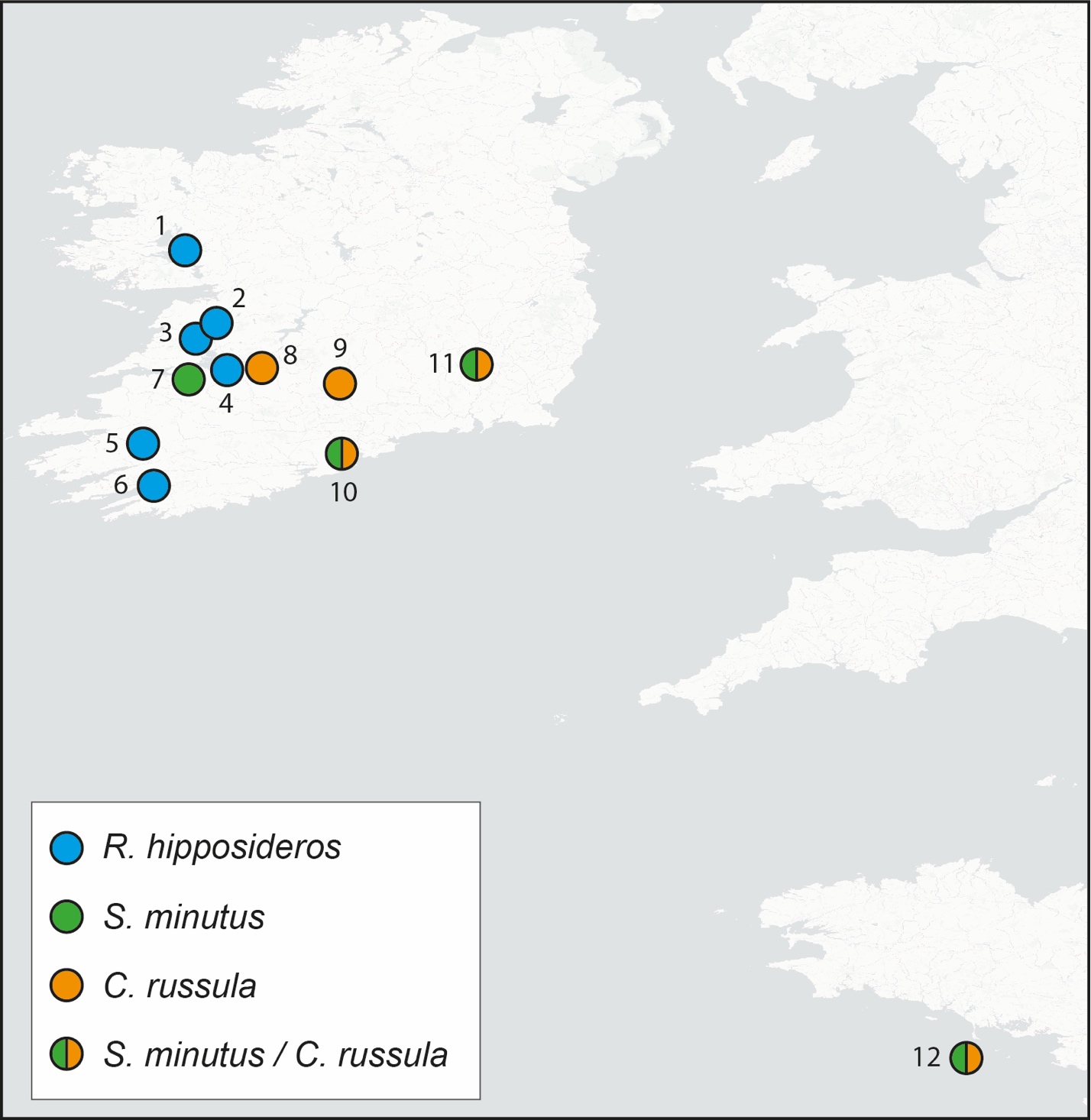

Figure S1. Sampling locations (numbered 1-11) in Ireland and Belle Île (12), France for lesser horseshoe bats (*Rhinolophus hipposideros*) in blue, pygmy shrews (*Sorex minutus*) in green and greater white-toothed shrews (*Crocidura russula*) in orange. See Table S1 for further details of numbered locations.

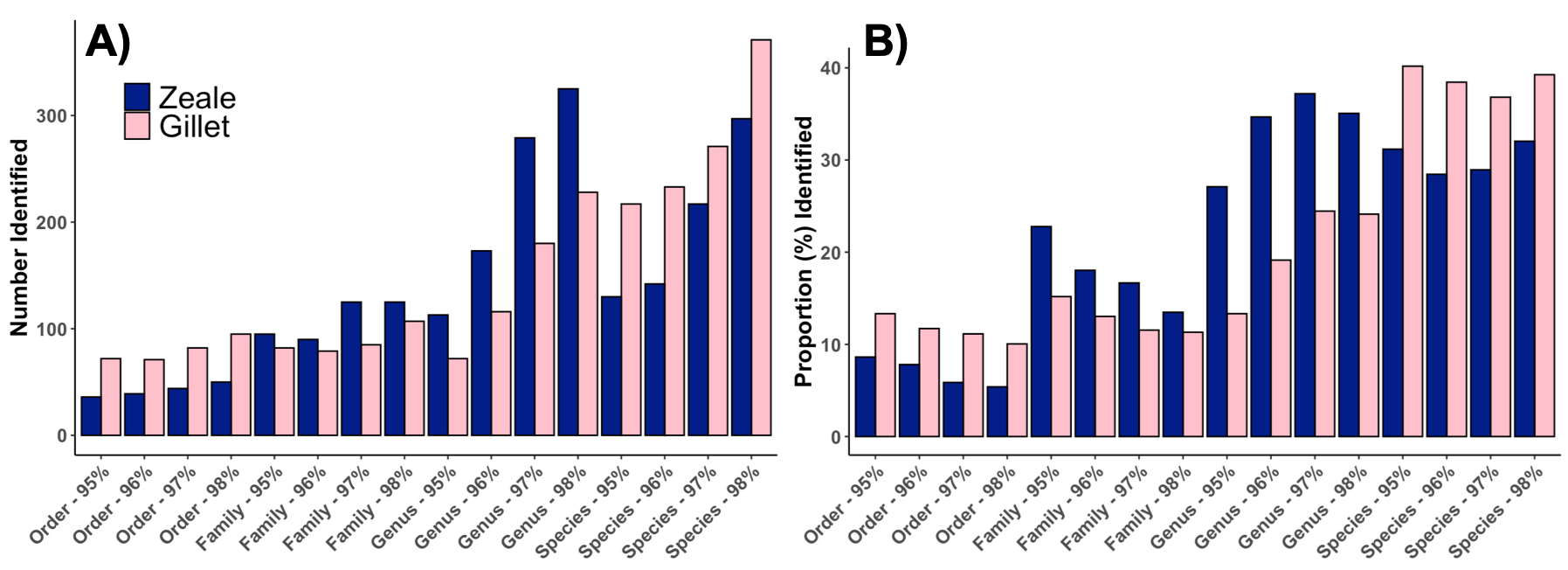

Figure S2. A) Number of MOTUs accurately assigned to order, family, genus and species level using different sequence clustering thresholds in sumaclust. B) Proportion of all MOTUs accurately assigned to order, family, genus and species level using different sequence clustering thresholds in sumaclust.

References

Alberdi A, Aizpurua O, Gilbert MTP, Bohmann K (2018) Scrutinizing key steps for reliable metabarcoding of environmental samples. Methods in Ecology and Evolution 9(1):134–147. doi: 10.1111/2041-210X.12849

Alberdi A, Gilbert MTP (2019) A guide to the application of Hill numbers to DNA ‐ based diversity analyses. Molecular Ecology Resources 19:804–817. doi: 10.1111/1755-0998.13014.

Andrews S (2010) FastQC: a quality control tool for high throughput sequence data.

Arrizabalaga-Escudero A, Clare EL, Salsamendi E, Alberdi A, Garin I, Aihartza J, Goiti U (2018) Assessing niche partitioning of co-occurring sibling bat species by DNA metabarcoding. Molecular Ecology 27(5):1273–1283. doi: 10.1111/mec.14508.

Boyer F, Mercier C, Bonin A, Le Bras Y, Taberlet P, Coissac E (2016) obitools: A unix-inspired software package for DNA metabarcoding. Molecular Ecology Resources 16(1):176–182. doi: 10.1111/1755-0998.12428.

Clare EL, Barber BR, Sweeney BW, Herbert PDN, Fenton MB (2011) Eating local: Influences of habitat on the diet of little brown bats (*Myotis lucifigus*). Molecular Ecology 20(8):1772-1780. doi: 10.1111/j.1365-294X.2011.05040.x.

Clare EL, Symondson WOC, Fenton MB (2014) An inordinate fondness for beetles? Variation in seasonal dietary preferences of night-roosting big brown bats (*Eptesicus fuscus*). Molecular Ecology 23(15):3633–3647. doi: 10.1111/mec.12519.

Edgar RC, Haas BJ, Clemente JC, Quince C, Knight R (2011) UCHIME improves sensitivity and speed of chimera detection. Bioinformatics 27(16):2194–2200. doi: 10.1093/bioinformatics/btr381.

Meiklejohn KA, Damaso N, Robertson JM (2019) Assessment of BOLD and GenBank – Their accuracy and reliability for the identification of biological materials. Plos One, 14(6):e0217084.

Oksanen J, Guillaume Blanchet F, Friendly M, Kindt R, Legendre P, McGlinn D, Minchin PR, O'Hara RB, Simpson GL, Solymos P, Henry M, Stevens H, Szoecs E, Wagner H (2019) vegan: Community Ecology Package. R package version 2.5-6.

Razgour O, Clare EL, Zeale MRK, Hanmer J, Schnell IB, Rasmussen M, Gilbert TP, Jones G (2011) High-throughput sequencing offers insight into mechanisms of resource partitioning in cryptic bat species. Ecology and Evolution 1(4):556–570. doi: 10.1002/ece3.49.

Rognes T, Flouri T, Nichols B, Quince C, Mahé F (2016) VSEARCH: a versatile open source tool for metagenomics. PeerJ 4:e2584. doi: 10.7717/peerj.2584.
